# The structure of enteric human adenovirus 41 - a leading cause of diarrhea in children

**DOI:** 10.1101/2020.07.01.181735

**Authors:** K. Rafie, A. Lenman, J. Fuchs, A. Rajan, N. Arnberg, L.-A. Carlson

**Author notes:** These authors contributed equally (K. Rafie, A. Lenman).

## Abstract

Human adenovirus (HAdV) types F40 and F41 are a prominent cause of diarrhea and diarrhea-associated mortality in young children worldwide. These enteric HAdVs differ strikingly in tissue tropism and pathogenicity from respiratory and ocular adenoviruses, but the structural basis for this divergence has been unknown. Here we present the first structure of an enteric HAdV - HAdV-F41 - determined by cryo-EM to a resolution of 3.8Å. The structure reveals extensive alterations to the virion exterior as compared to non-enteric HAdVs, including a unique arrangement of capsid protein IX. The structure also provides new insights into conserved aspects of HAdV architecture such as a proposed location of protein V, which links the viral DNA to the capsid, and assembly-induced conformational changes in the penton base protein. Our findings provide the structural basis for adaptation to a fundamentally different tissue tropism of enteric HAdVs.

## MAIN TEXT

### Introduction

Adenoviruses (AdVs) are common pathogens in human, causing diseases in airways, eyes, and intestine, but also in the liver, urinary tract, and/or adenoids(1). To date, more than 100 human adenovirus (HAdV) types have been isolated, characterized, and classified into seven species (A-G)(2). In addition to causing serious diseases in humans, several adenoviruses are being explored as vaccine vehicles against infectious diseases such as COVID-19, MERS, Ebola disease, AIDS, Lassa fever, and Zika disease(3). This includes a vaccine candidate based on HAdV-F41 which elicited neutralizing antibodies against MERS-CoV *in vivo*(4). The two sole members of HAdV species F, HAdV-F40 and HAdV-F41, stand out as the only HAdVs with a pronounced gastrointestinal tropism. These so-called enteric adenoviruses are a leading cause of diarrhea and diarrhea-associated mortality(5) in young children, inferior only to Shigella and rotavirus(6). Diarrhea is estimated to cause ∼530,000 deaths/year in children younger than five years worldwide(7). Thus, there are strong incentives to understand the structural and molecular basis of enteric HAdVs.

AdVs are double-stranded DNA viruses with an approximately ∼35 kbp genome sheltered in a large (∼950 Å in diameter), non-enveloped capsid with icosahedral symmetry(8–13). At each of the twelve capsid vertices, penton base subunits organize as homopentamers(14), anchoring the N-terminal tails of the protruding, trimeric fibers(15, 16) to the capsid. Another so-called major capsid protein is the hexon protein(17), present in 240 trimers per virion. Hexons are the main structural component of the virion facets and are organized to give the virion a pseudo *T=25* symmetry (Fig. 1A). Hexon assemblies are stabilized by minor capsid protein IIIa (pIIIa), pVI and pVIII (located in the capsid interior) and by pIX (exposed on the capsid exterior)(9, 10, 18). To date, two human adenoviruses, HAdV-C5(11, 12, 19, 20) and HAdV-D26(21), but also individual capsid proteins or their subdomains(14, 17, 22) of multiple adenovirus types, have been structurally determined at high resolution.

**Figure 1:**
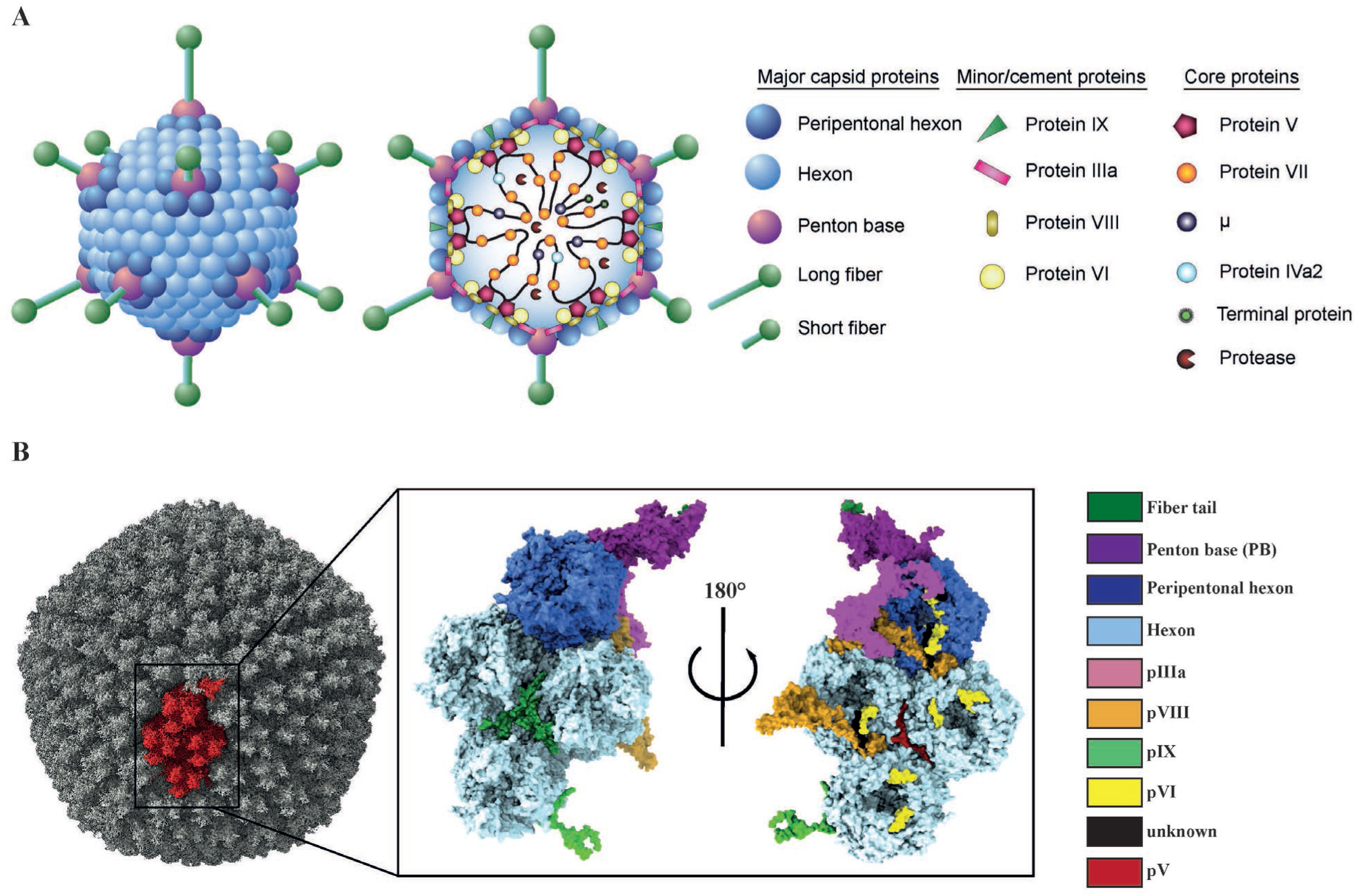
The overall structure of HAdV-F41. (A) Schematic representation of the capsid and core structure of HAdV-F41. (B) Surface representation of the HAdV-F41 electron density with one asymmetric unit highlighted in red (left) and a surface representation of the asymmetric unit of the HAdV-F41 atomic model viewed from the virion exterior (middle) and interior (right).

The enteric HAdVs have adapted to a distinct tissue tropism from other adenoviruses, which is presumably reflected in their capsid structure. One major known difference is that enteric HAdVs contain two different types of fiber proteins, long and short(23, 24), whereas other AdVs contain only one type of fiber. Virions dock on to cells through fiber interactions with cellular receptors, followed by internalization mediated by penton base interactions with cellular integrins(25). All other HAdVs contain a conserved, integrin-interacting Arg-Gly-Asp (RGD) motif in the penton base(26). Strikingly, the enteric HAdV-F40 and -F41 lack this conserved RGD motif and thus use different integrins for entry(27), which may explain their different and much slower entry mechanism(28, 29). Knowledge about the capsid proteins, their structural organization and host molecule interactions is important for design and development of adenoviruses as vectors and vaccine vehicles.

Despite the medical importance of enteric adenoviruses as a major cause of childhood mortality through diarrhea, the structural basis for their infection is not known. We used cryo-EM to determine near-atomic structures of the HAdV-F41 virion at pH=7.4 and at pH=4.0, the latter set as an average of the diurnal pH in the stomach of young children(30). These structures reveal this enteric adenovirus as having a pH-insensitive capsid with extensive surface remodeling as compared to non-enteric HAdVs. We further propose a conserved location of core protein V (pV), which links the adenovirus genome to the capsid. Lastly, we describe the assembly-induced structural changes to the penton base protein. We believe that these findings will lay the foundation for a detailed molecular understanding of enteric adenoviruses, how to prevent their infection, and how to explore AdVs as vehicles for vaccine development.

### Results

#### The structure of HAdV-F41 reveals a pH-resistant capsid with an altered surface charge distribution

To elucidate the structural basis of enteric adenovirus infection, we determined the structure of HAdV-F41 using cryo-EM. In parallel, the genome of the purified virus (strain “Tak”) was sequenced, revealing one protein-coding mutation (Val77Ala in protein VIII) compared to the deposited sequence for the same strain (GenBank: DQ315364.2). Further, its proteome was determined using the high-recovery filter-aided sample preparation (FASP) mass spectrometry(31), revealing a total of 22 viral proteins present in the purified virus (Supplementary Table 1). At an average resolution of 3.8 Å, the three-dimensional reconstruction of HAdV-F41 had continuous electron density with well-defined secondary structure elements and side-chain density (Supplementary Figure 1, Supplementary Movie 1). Local resolution estimates revealed a resolution of better than 3 Å for large parts of the icosahedral capsid (Supplementary Figure 1). This allowed us to build and refine an atomic model of the asymmetric unit (ASU), which describes the icosahedral part of the virus (Fig. 1B, Supplementary Movie 2). The final ASU model contained four hexon homotrimers, single chains of the penton base protein and pIIIa, ten chains of pVI, two chains of pVIII, three chains of the triskelion protein IX, and five chains of unknown identity, revealing known and unknown protein-protein interaction surfaces (Supplementary Table 2). Electron density for the fibers was present at the interface with the icosahedral capsid but was of insufficient quality for extensive model building due to increasing flexibility in more distal parts. Compared to the two other reported structures of human adenoviruses, HAdV-C5 (PDB: 6B1T(20)) and HAdV-D26 (PDB: 5TX1(21)), the sequence identity of the capsid proteins ranges from 30-80% (Supplementary Table 3). This generally correlated with the average structural similarity between the three structures, with more divergent proteins showing higher structural difference in terms of Ca root mean square deviation (RMSD) (Supplementary Table 3).

In their adaptation to gastrointestinal tropism, a major obstacle for enteric adenoviruses was likely the passage through the low pH of the stomach to their intestinal site of infection. To investigate the adaptation to the hostile environment of the stomach, we solved the structure of HAdV-F41 at pH=4.0, which resembles the diurnal average gastric pH of young children(30) (Fig. 2A). At a resolution of 5.0 Å the overall structure of the capsid at pH=4.0 was largely unchanged (overall Ca RMSD 3.0 Å for proteins listed in Fig. 2B) and no drastic local movements were observed (Fig. 2B), showing the resistance of the enteric adenovirus capsid to the gastric pH. We reasoned that the gastrointestinal adaptation might also have altered the distribution of acidic and basic residues exposed on the outer surface of the capsid. To investigate this, the surface charge distribution of HAdV-F41 along with the other two existing human adenovirus structures (HAdV-C5 and HAdV-D26) were calculated for pH=7.4. A visual comparison of the charge distribution revealed significant differences between the three HAdVs (Fig. 2C). The exposed surface of HAdV-D26 is almost entirely covered by negative charge at pH=7.4, and HAdV-C5 has mostly negatively charged surfaces on the top of the hexons (in a calculation underestimating the amount of negative charge due to exclusion of the flexible, and highly negatively charged, hypervariable region 1, which was not built in any of the HAdV-C5 structures(19, 20, 32–35)). By comparison, the capsid of HAdV-F41 is predominantly uncharged at pH=7.4 (Fig. 2C). The surface charge distribution of HAdV-F41 at pH=4.0 revealed two distinct regions as still being relatively uncharged at this extreme pH: the N-terminal part of pIX, largely occluded between hexons, and the solvent-exposed loops at the top of the hexons (Fig. 2C).

**Figure 2:**
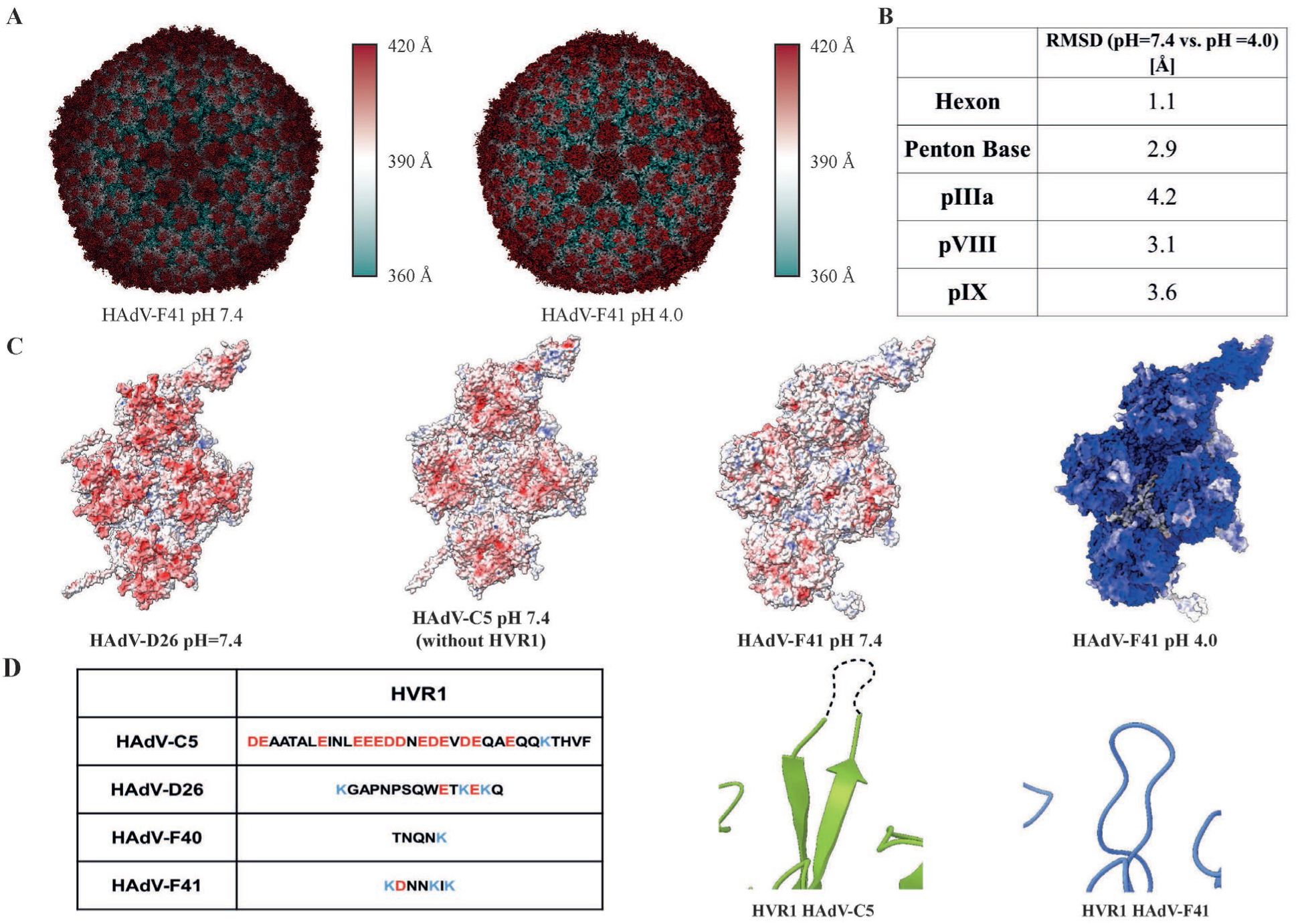
Structure and surface charge distribution of HAdV-F41 at pH=7.4 and pH=4.0. (A) Surface representation of the HAdV-F41 electron density at pH=7.4 and pH=4.0, colored by distance from the virion center. (B) pH-dependent structural changes in selected HAdV-F41 capsid proteins as measured by RMSD at Cα level. (C) Surface charge distribution for the atomic models of HAdV-C5 (PDBID: 6CGV), -D26 (PDBID: 5TX1) and -F41 at pH=7.4 as well as HAdV-F41 at pH=4.0. Red represents a local net negative charge, blue positive, and white uncharged. (D) Comparison of the HVR1 sequences between HAdV-C5, -D26, -F40 and -F41. (E) Cartoon representation of the HVR1-containing loop for HAdV-C5 and HAdV-F41. The unbuilt loop for HAdV-C5 is shown as a black dashed line.

Whereas the overall structural fold of hexon chains is conserved among adenoviruses, they differ in the seven hypervariable regions (HVRs). Comparing HAdV-F41 (at pH=7.4) to HAdV-C5 and HAdV-D26, substantial differences were found in all seven HVRs (Supplementary Table 5). In particular, HVR1 stands out in the comparison since it is much shorter in enteric HAdVs (Fig. 2D).

The absence of the highly negatively charged HVR1 loop in all HAdV-C5 structures(11, 20) indicates flexibility. On the other hand, the near-eliminated HVR1 in HAdV-F41 is a short loop with a rigid conformation that allowed tracing the entire length of the polypeptide (Fig. 2E).

Taken together, these findings show that the capsid of enteric HAdV-F41 is structurally unperturbed by exposure to stomach-like pH and has evolved to expose fewer charged residues on its exterior as compared to non-enteric HAdVs, most prominently exemplified by a near-complete deletion of the hypervariable region 1.

#### Protein IX arranges in a unique manner in HAdV-F41

Among the so-called minor capsid proteins, pIX forms the most extended and complex arrangements. In previously reported structures of HAdV-C5 and HAdV-D26, this amounts to a tight mesh of ordered protein density that stretches through the canyons between hexons across the virion surface(11, 21, 36) (Fig. 3A). In HAdV-F41, only the N-terminal residues 1-58 (henceforth ‘pIX-N’) were sufficiently ordered to trace the protein chain, thus ruling out the same sort of ordered, virus-spanning pIX cage seen in other HAdVs (Fig. 3A). Analogously to other HAdVs, pIX-N trimerizes to form a triskelion (Fig. 3B) located between three non-peripentonal hexon units (Fig. 1B). Each facet of the virion harbors four copies of pIX-N triskelion in two distinct structural surroundings: three copies at the local 3-fold (L3) symmetry axes of each asymmetric unit, and one at the icosahedral 3-fold (I3) symmetry axis at the center of the facet (Supplementary Figure 2). The conformations of pIX-N in these two surroundings are virtually identical (Fig. 3B). The three pIX chains come together at the center of this triskelion, where interactions between residues Phe12, Phe17 and Tyr20 from each chain form a hydrophobic core (Fig. 3C). This hydrophobic core is differently arranged compared to the non-enteric HAdV-C5 and HAdV-D26 (Supplementary Figure 2) and has more large hydrophobic residues at its center.

**Figure 3:**
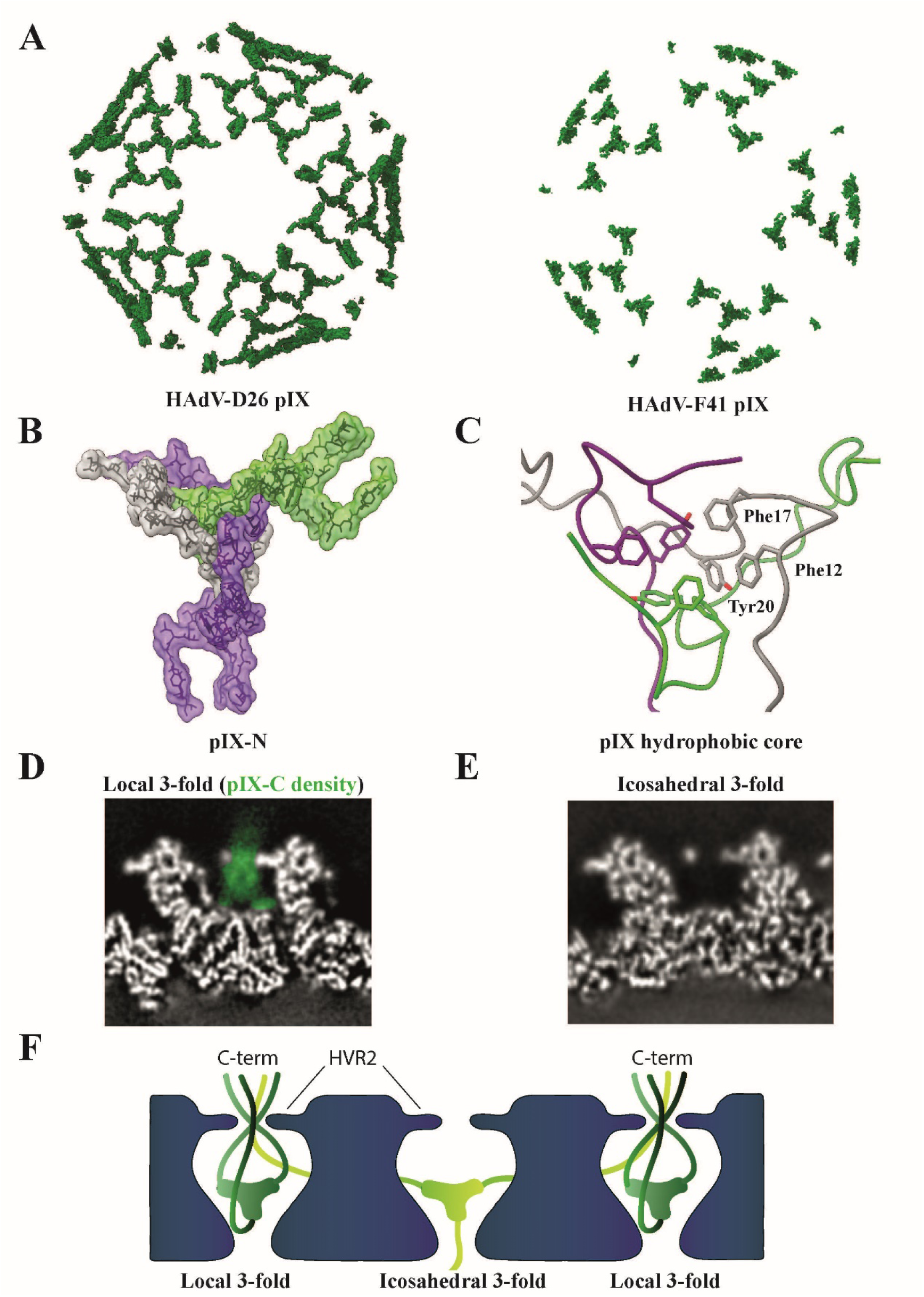
The triskelion-forming protein IX assembles in a unique way in HAdV-F41. (A) Surface representation of pIX assembly in HAdV-D26 and HAdV-F41 including all ordered protein density. (B) Graphical representation of HAdV-F41 pIX-N triskelion assembly. The three pIX chain atoms (green, grey, purple) are shown in stick representation covered by a semitransparent surface. (C) Close-up view of the hydrophobic core at the center of the pIX triskelion assembly, formed by residues Phe12, Phe17 and Tyr20 of each pIX chain. (D) Computational slice through the electron density of HAdV-F41 at pH=7.4, at the position of pIX at the local 3-fold axis. Electron density is white except the proposed density of pIX-C which is highlighted in green. (E) As (D), but at the icosahedral 3-fold axis. (F) Schematic representation of one possible arrangement of pIX-C.

Sequence comparisons of pIX revealed that pIX-N has a higher sequence homology to HAdV-C5 and HAdV-D26 than its C-terminal part (residues 59-133, ‘pIX-C’) (Supplementary Figure 2). However, the sequences of pIX-C are near-identical in HAdV-F41 and the related HAdV-F40, indicating conservation between enteric adenoviruses. Mass spectrometry analysis detected the entire pIX sequence in the purified HAdV-F41, confirming the presence of pIX-C in the purified virions (Supplementary Table 1). We reasoned that the substantial protein mass corresponding to pIX-C, emanating in the constricted space between the hexons, should be visible in the electron density map at a lower threshold even if it is flexible. Indeed, the interhexonal space above pIX-N harbors electron density corresponding to a flexible protein at the local 3-fold axes (Fig. 3D, Supplementary Figure 3). This density appears to pass to the outside of the capsid between the HVR2 loops of the three surrounding hexons. In a localized asymmetric reconstruction(37), the three HVR2 could be resolved in their entirety (Supplementary Figure 4), showing that they are present in a single conformation and form a constriction of defined size (Supplementary Figure 3) from which the C-termini of pIX protrude. In contrast, there is clearly no electron density above pIX-N at the icosahedral 3-fold position (Fig. 3E), showing that the spatial organization of pIX-C differs between these two positions (Fig. 3F). Taken together, these data reveal a very different arrangement of pIX in enteric adenoviruses as compared to respiratory (C5) and ocular (D26) HAdVs. In HAdV-F41, the C-terminal half of pIX is flexible and exposes it’s C-terminus to the capsid exterior at three of the four pIX positions in each virion facet.

#### The HAdV-F41 penton base undergoes assembly-induced conformational changes

Located on the five-fold symmetry axes of adenovirus capsids, the penton base (PB) protein forms a homopentamer that contains integrin-binding motifs and serves as an assembly hub connecting the icosahedral capsid to the fibers (Fig. 1A). In our reconstruction of the entire HAdV-F41, the electron density for the PB was less well resolved than other parts of the capsid. To improve the map of the PB, we performed a localized asymmetric reconstruction of the PB monomer (Supplementary Figure 4). The improved map allowed the building of an atomic model for the PB, which was placed into a composite atomic model of the entire asymmetric unit. Overall, the HAdV-F41 PB is very similar to the PB in HAdV-C5(11, 12, 20) and HAdV-D26(21), with a β-sheet rich fold that can roughly be divided into four domains: crown, head, body and tail, with the body and head as the main domains, separated by loop regions (Fig. 4A). During assembly of the virus particle, the PB homopentamer forms a plethora of interactions with peripentonal hexons, the fiber, and the minor capsid protein pIIIa (Supplementary Table 2). To investigate conformational changes induced during the PB assembly process, we solved the structure of a recombinantly expressed HAdV-F41 PB homopentamer in solution (free PB -fPB) by cryo-EM. The map had an average resolution of 3.7 Å (Supplementary Figure 5), allowing for the placement of an atomic model (Fig. 4B). Comparing the atomic models of the fPB and the virion-bound PB (vPB), the overall Cα-shift (RMSD) was very small (∼0.9 Å). However, color-coding vPB by its structural deviations from fPB revealed regions with higher RMSD, indicating localized assembly-induced conformational changes (Fig. 4C, Supplementary Movie 3). Moreover, four sequence segments that were built in the vPB model could not be built in the fPB model (Fig. 4A), indicating that these regions are disordered in solution and only become stabilized in a defined conformation upon assembly into the virion. One such region is the tail, a 17 residue (Thr33-Gly49) random coil region (Fig. 4A), which is disordered in the fPB (Fig. 4B). It is stabilized through interactions with two loop regions from pIIIa (Supplementary Table 2). The sequence of the tail domain is largely conserved between HAdVs (Supplementary Figure 6), suggesting a conserved role as an assembly motif. The second motif (Tyr419-Leu429) becoming ordered upon assembly is an α-helix consisting of residues Gln416-Thr427, located close to the five-fold axis of the penton base (Fig. 4D), and close to where the fiber binds. Although the HAdV-F41 capsid map shows only weak density for the proximal fiber in connection with the PB, the vPB structure did allow tracing of a fragment of the conserved fiber tail (Supplementary Figure 6, Supplementary Table 3). Thus, the folding of Gln416-Thr427 may be dependent upon interactions within the capsid and/or binding of the fiber to the PB. Additional assembly-dependent interactions take place in two loop regions located between residues Val70-Asn110 in the body domain (Fig. 4D). These loops are disordered in the fPB but well resolved in the vPB where their conformation is stabilized by interactions with the peripentonal hexon. The first loop (Ser74-Ser79) - located at the top of the body - is stabilized as an extended coil structure upon interaction with hexon chain Ser663-Tyr671 loop. The second region (Thr100-Gln107) - located at the bottom of body - is stabilized as a short α-helix upon binding to an uncharged pocket formed by peripentonal hexon residues Ala623-Ile640.

**Figure 4:**
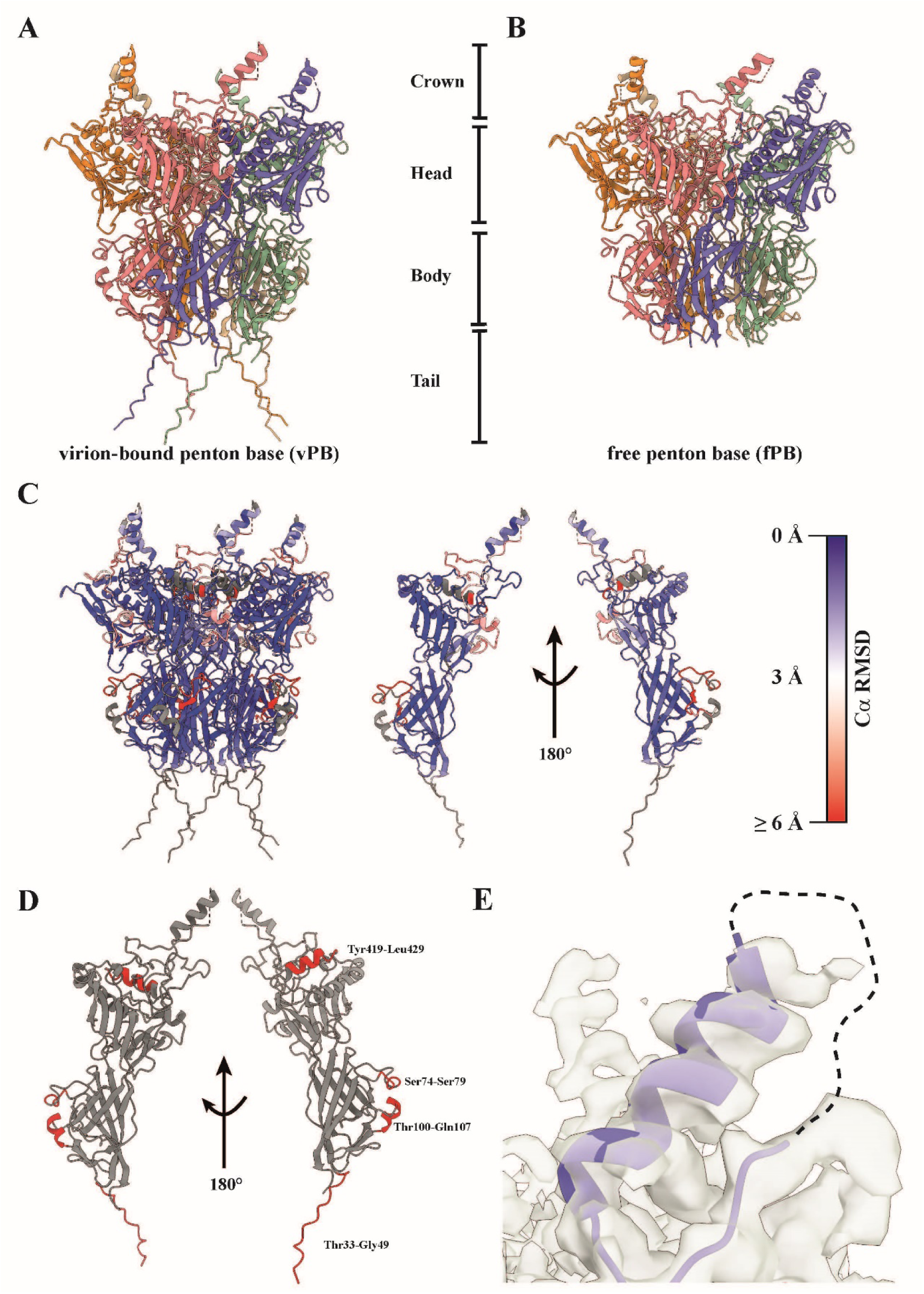
The HAdV-F41 penton base undergoes assembly-induced conformational changes. (A) Cartoon representation of the virion-bound penton base (vPB), which can be divided into a crown, head, body and tail. (B) Cartoon representation of the free penton base (fPB). (C) Cartoon representation of the vPB and a single vPB monomer chain, each colored by Ca root-mean-square-deviation (RMSD) indicating the local degree of difference in Ca positioning between the vPB and fPB structures. (D) Cartoon representation of a single vPB monomer chain (grey). Missing residues in the fPB structure are highlighted in red. (E) Cartoon representation of the helix and disordered loop region containing the integrin-binding IGDD motif (dashed line) located in the crown. The electron density is shown as a transparent surface.

Peculiarly for enteric HAdVs, the otherwise conserved integrin-binding RGD motif has been replaced by IGDD in HAdV-F41 (RGAD for HAdV-F40). In the HAdV-F41 structure the IGDD-containing loop is the only surface-exposed part of the PB for which we find no continuous electron density (Fig. 4E), despite being among the shortest loop of all HAdVs(26) (Supplementary Figure 6). This parallels the observed flexibility of the RGD motifs in the two previously reported structures of HAdVs(19–21), indicating that the function of the IGDD sequence may be dependent on it being flexible until interacting with a target molecule.

In summary, a comparison of the PB pentamer in solution and in the virus capsid revealed several distinct motifs that become folded only upon assembly of the PB into the capsid, and further revealed that the non-canonical integrin-binding motif IGDD is disordered also in the context of the assembled virus.

#### The DNA-binding protein V is located at a conserved position at the inner face of the capsid

After initial model building of the virion at pH=7.4, the asymmetric unit contained five peptide chains that still had not been assigned an identity. Four of the five chains were deemed too short to assign an identity to. Three of those four chains interlace with different copies of pVI at positions where pVI interacts with pXIII or pIIIa at the inner face of the capsid (Supplementary Figure 7, Supplementary Movie 2). The fifth unidentified peptide chain is markedly longer and is located at the inner face of the capsid, in a pocket formed by three non-peripentonal hexons (Fig. 5A-B, Supplementary Table 6). Its electron density is resolved well enough to identify large side-chain residues and we thus reasoned that a structural bioinformatics workflow might be devised to reveal its identity. With the initial constraint being only that residues number 4 and 21 in the identified 24mer peptide must have large side chains, we utilized a combination of the proteomics data, exclusion of proteins with known locations, real-space refinement scores and other considerations to elucidate its identity (Supplementary Methods 1, Supplementary Figure 8). After sequential exclusion of candidates based on these criteria, a single most likely candidate remained, a sequence from the center of protein V (pV): Gln170-Asp194. pV has been reported to bind directly to DNA and to pVI, thereby bridging the core and the surrounding capsid, but it has not been localized in any adenovirus structure. The built sequence of pV fits the density without any clashes or unlikely interactions with surrounding proteins, and furthermore shows a high degree of sequence conservation with pV in HAdV-C5 and HAdV-D26 (Fig. 5C, D, Supplementary Movie 4). In both hitherto published HAdV structures, HAdV-C5(20) and HAdV-D26(21), there is similarly shaped electron density at the corresponding position of the virus capsid (Fig. 5E). In HAdV-C5 no atomic model was built into it(20). Similarly, at the corresponding position in the HAdV-D26 structure, two shorter peptide chains of unknown identity were placed(21). The location of the 24mer chain of pV is such that both the N-and C-terminus of pV, which are not resolved in our structure, may protrude towards the interior of the virion in agreement with the proposed role of pV to link the viral genome to the capsid. In summary, we used a systematic structural bioinformatics approach to propose a likely conserved position of pV in human adenoviruses.

**Figure 5:**
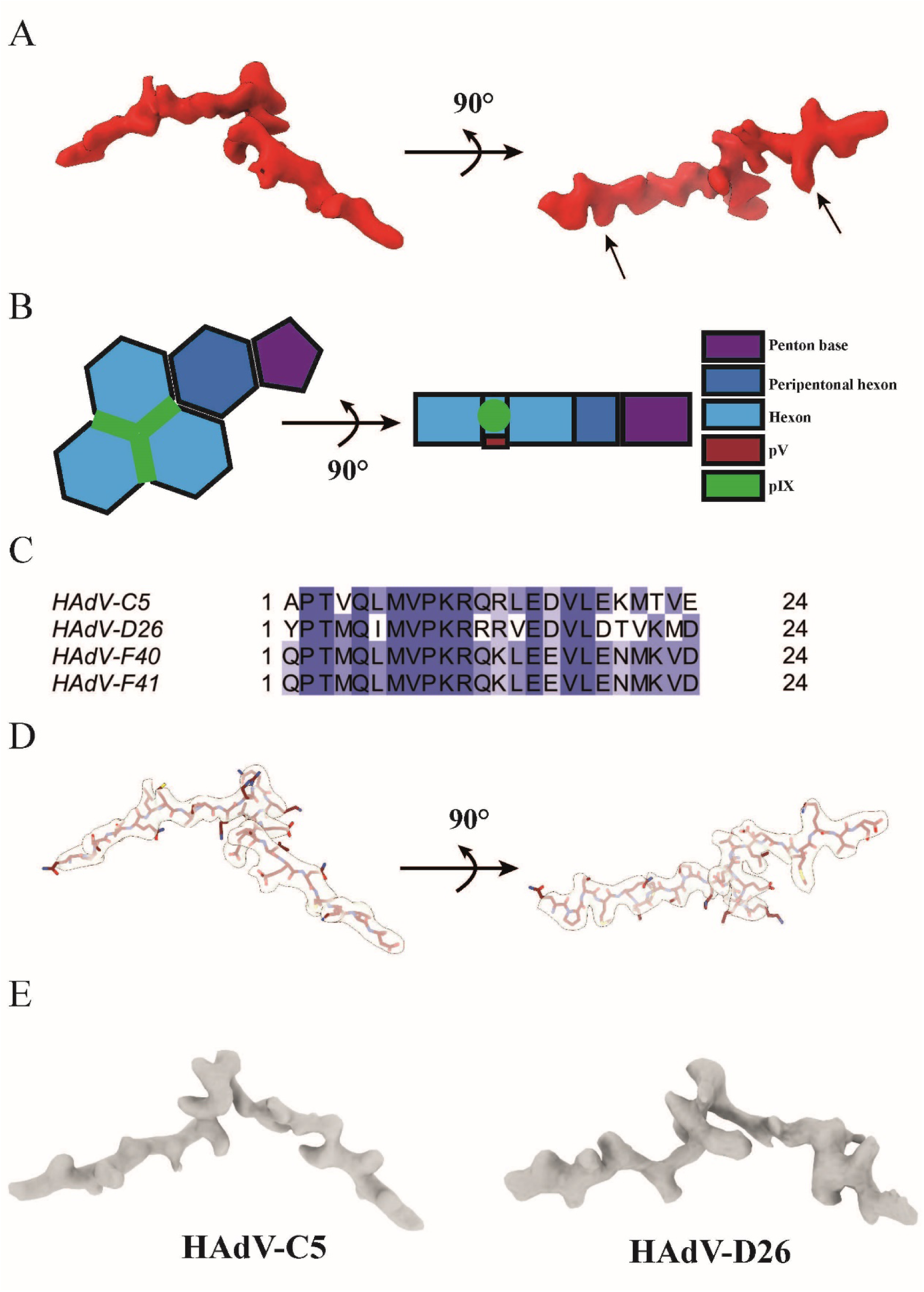
Location of DNA-binding protein V at the interface of the three non-peripentonal hexon subunits. (A) Surface representation of pV electron density. Arrows indicate the positions fixed during bioinformatics analysis. (B) Schematic representation of pV and its location in the HAdV-F41 asymmetric unit. (C) Alignment of the identified pV amino acid sequence from HAdV-C5, -D26, -F40 and -F41. Coloring represents % sequence identity, with dark blue illustrating 100% homology. (D) Graphical representation of the modelled HAdV-F41 pV peptide, shown in maroon stick representation covered by the corresponding electron density, shown as transparent surface. (E) Surface representation of the HAdV-C5 (EMD-7034(20)) and HAdV-D26 (EMD-8471(21)) electron densities located at the same position in their respective asymmetric units.

### Discussion

Here, we present the structure of a major cause of diarrhea and diarrhea-associated mortality in children: the enteric human adenovirus HAdV-F41. As the first structure of an adenovirus with pronounced gastrointestinal tropism, it reveals a capsid stable to stomach pH, with substantial changes to the virion surface as compared to respiratory and ocular HAdVs. Overall, HAdV-F41 has fewer charged, i.e. pH-dependent, residues exposed on the surface of its capsid. This is especially prominent at the top of the hexons where HVR1 is long, and rich in negatively charged residues in HAdV-C5. This allows interaction of HAdV-C5 with lactoferricin through a charge-dependent mechanism(35), which contributes to an extended tissue tropism(39, 40). Evolution of HAdV-F41 has resulted in a largely truncated and less charged HVR1 (Fig. 2D), seemingly to adapt to the specific conditions in the gastrointestinal tract. Another major change to the capsid exterior of HAdV-F41 is the starkly different conformation of protein IX (pIX). Instead of forming a virus-covering, rigid mesh, the C-terminal half of pIX (pIX-C) is flexible and protrudes to the outside of the capsid, kept in place by hexon HVR2-containing loops. Strikingly, pIX-C density is observed above all pIX-N trimers except at the icosahedral 3-fold axis (Fig. 3D-E, Supplementary Figure 2). In principle, each pIX-C strand could either emerge in *cis* (i.e. above the same pIX-N trimer to which it belongs), or stretch across the virion surface to emerge above another pIX-N trimer (in a *trans* arrangement). The length of pIX-C in HAdV-F41 is compatible with both of these arrangements. A *cis* arrangement of all strands would be reminiscent of pIX in some non-human adenoviruses, in which the conformation of pIX-C is also more defined(36, 41, 42). Whereas our data don’t allow tracing of individual strands of pIX-C, the lack of any pIX-C density above the pIX-N trimer at the icosahedral 3-fold rules out such a pure *cis* arrangement of pIX-C. One possible, parsimonious interpretation would be that the central pIX trimer adopts a *trans* arrangement, donating one pIX-C strand to each of the three pIX trimers at local 3-fold positions which in turn have their pIX-Cs in *cis* (Fig. 3F). Whether this model is correct or not, it is clear from our data that pIX arranges in a unique manner in enteric adenoviruses compared to other HAdVs studied to date. All these modifications to the virion surface of HAdV-41 are likely related to the different set of interactors, and different pH range that this virus encounters throughout the gastrointestinal tract. Other gastrointestinal viruses interact with components such as bile (calicivirus)(43) and lipopolysaccharides (poliovirus(44), mouse mammary tumor virus(45)), which is crucial for infection of these viruses. Besides low-pH resistant interactions of HAdV-F41 with gastrointestinal phospholipids(46), little is known about the HAdV-F41:gastrointestinal interactome. Finding such interaction partners, e.g. of the disordered and exposed pIX-C region, will yield further insights into the infection cycle and tropism of enteric adenoviruses. Our study further unveiled how several motifs in the HAdV-F41 PB are disordered in solution and only adopt a defined conformation upon assembly into the virus capsid, laying out another piece of the still largely unfinished puzzle of adenovirus assembly(10). The observation that the modified integrin-interacting motif of the HAdV-F41 PB is still disordered in the assembled virus particle highlights the need for structural studies of the interactions with its proposed binding partner, laminin-binding integrins(27). Biochemical data have defined protein V (pV) as a key protein linking the adenovirus genome to the capsid(47), but in spite of its conserved function in adenoviruses it had not been located in the adenovirus capsid. Here we propose the point of interaction of pV to the interior of the capsid, and provide data suggesting that this position, at the junction between three non-peripentonal hexons, is conserved between HAdVs. Previous biochemical data have not suggested pV to interact with the hexons, but have instead suggested interactions between pV and the minor protein pVI(47, 48). These data are not mutually exclusive with our identification of the pV anchoring point to the hexons, since most of the pV sequence is still unaccounted for in the structure, and several copies of pVI are found in the vicinity of the anchoring point where they may form additional interactions. Taken together, the structure of the enteric adenovirus HAdV-F41 revealed key conserved aspects of adenovirus architecture as well as highly divergent features of enteric adenoviruses, thus laying the foundation for structure-based approaches to preventing this prominent cause of diarrhea-associated mortality in young children, and, for further development of these structurally divergent AdV types as vaccine vehicles.

### Materials and Methods Virus propagation and purification

Human A549 cells (kind gift from Dr. Alistair Kidd) were maintained in Dulbecco’s modified Eagle medium (DMEM; Sigma-Aldrich) supplemented with 5% fetal bovine serum (FBS; HyClone, GE Healthcare), 20 mM HEPES (Sigma-Aldrich) and 20 U/ml penicillin + 20 μg/ml streptomycin (Gibco).

For HAdV-F41 (strain Tak) propagation, 30 bottles of A549 cells (175cm^2^, at a 90% confluency) were infected with HAdV-F41 inoculation material (produced in A549 cells) in 5 ml of growth media (1% FBS) for 90 min on a rocking table at 37°C. Thereafter additional 25ml of growth media (1% FBS) was added to each flask and the cells were further incubated at 37°C. Infected cells were harvested after approximately one week, or when cells displayed clear signs of cytopathic effect. Cells were collected by centrifugation, resuspended in DMEM and disrupted to release virions by freeze thawing and by addition of equal volume of Vertrel XF (Sigma-Aldrich). After vigorous resuspension, the cell extract was centrifuged at 3000 rpm for 10 min. The upper phase was transferred onto a discontinuous CsCl gradient (densities: 1.27 g/mL, 1.32 g/mL, and 1.37 g/mL, in 20 mM Tris-HCl, pH=8.0; Sigma-Aldrich) and centrifuged at 25000rpm in a Beckman SW40 rotor for 2.5h at 4°C. The virion band was collected and desalted on a NAP column (GE Healthcare) into sterile PBS.

### Protein Identification by mass spectrometry

#### Protein digestion

The samples were split for tryptic and chymotryptic digestion and processed using a modified protocol of filter-aided sample preparation (FASP) (31). In brief, triethylammonium bicarbonate (TEAB) was added to a final concentrations of 50 mM TEAB prior to reduction using 100 mM dithiothreitol at 56°C for 30 min. The reduced samples were loaded onto 10 kDA MWCO Pall Nanosep centrifugal filters (Sigma-Aldrich), washed with 8M Urea, 1% sodium deoxycholate (SDC) and alkylated with 10 mM methyl methane thiosulfonate. Two step digestion was performed on filters using trypsin and chymotrypsin as digestive enzymes in 50 mM TEAB, 0.5% SDC buffer. The first step was performed overnight and the second step, with an additional portion of proteases, for four hours the next day. Tryptic digestion was performed at 37°C using Pierce MS-Grade Trypsin Protease (Thermo Fisher Scientific). Chymotryptic digestion was performed at room-temperature using Pierce MS-Grade Chymotrypsin Protease (Thermo Fisher Scientific). The peptides were collected by centrifugation and SDC was precipitated by acidifying the sample with TFA (final concentration 1%). The digested sample was desalted using Pierce Peptide Desalting Spin columns (Thermo Scientific) according to the manufacturers protocol.

#### LC-MS/MS

The digested and desalted samples were analysed using a QExactive HF mass spectrometer interfaced with an Easy-nLC 1200 liquid chromatography system (both Thermo Fisher Scientific). Peptides were trapped on an Acclaim PepMap 100 C18 trap column (100 μm x 2 cm, particle size 5 μm, Thermo Fischer Scientific) and separated on an in-house packed analytical column (75 μm x 300 mm, particle size 3 μm, Reprosil-Pur C18, Dr. Maisch). A stepped gradient used was from 7% to 35% solvent B in 97 min followed by an increase to 48% in 8 min and to 100% solvent B in 5 min at a flowrate of 300 nL/min. Solvent A was 0.2% formic acid and solvent B was 80% acetonitrile in 0.2% formic acid. The mass spectrometer was operated in data-dependent mode (DDA) where the MS1 scans were acquired at a resolution of 60000 and a scan range from 400 to 1600 m/z. The 10 most intense ions with a charge state of 2 to 4 were isolated with an isolation window of 1.2 m/z and fragmented using normalized collision energy of 28. The MS2 scans were acquired at a resolution of 30000 and the dynamic exclusion time was set to 20 s.

#### Database search

Data analysis was performed using Proteome Discoverer (Version 1.4, Thermo Fisher Scientific). The data was searched against an in-house database containing the amino acid sequences of HAdV-F41. Mascot (Version 2.5.1, Matrix Science) was used as search engine with a precursor mass tolerance of 5 ppm for MS1 and 30 mmu for MS2 spectra. Tryptic peptides were accepted with a maximum of one missed cleavage, chymotryptic peptides with maximum three missed cleavages. Variable modification of methionine oxidation and fixed methylthio of cysteines were selected. The Mascot Significance Threshold for peptides was set to 0.01.

### Cryo-EM sample preparation

Purified HAdV-F41 was used at 4.2 mg/mL (pH=7.4) and 1.6 mg/mL (pH=4.0). The recombinant HAdV41 penton base (PB41) was purified as described before(27) and used at 1 mg/mL in PBS buffer, supplemented with 5% glycerol. A HAdV41 sample at pH=4.0 was prepared by adding 2 μL of a 0.5 M citric acid / 1 M Na2HPO4 pH=3.4 solution to 25 μL of HAdVF-41 pH=7.4 followed by incubating on ice for 15 minutes. Samples were vitrified on Quantifoil Cu R200 2/2 (Electron Microscopy Sciences, Cat#: Q2100CR2) and Quantifoil Cu R200 1.2/1.3 (Electron Microscopy

Sciences, Cat#: Q3100CR1.3) grids for the virus particles and the recombinant protein, respectively. Prior to sample application the grids were glow discharged using a Pelco easiGlow device (Ted Pella Inc.) at 15 mAmp for 30 s. Sample was applied by transferring 3 μL sample onto the glow-discharged side of the grid, blotted and plunge frozen in liquid ethane, using a Vitrobot plunge freezer (Thermo Fisher Scientific), with the following settings: 22 °C, 80% humidity, Blot-force = −20 and a blotting time of 3 s. For HAdVF-41 pH=7.4 sample was applied twice with a blotting step, using the same settings as above, between applications(49).

### Data collection

All data were collected on an FEI Titan Krios transmission electron microscope (Thermo Fisher Scientific) operated at 300 keV and equipped with a Gatan BioQuantum energy filter and a K2 direct electron detector. A condenser aperture of 70 μm (HAdV-F41 pH=7.4 & 4.0) and an objective aperture of 100 μm were chosen for data collection. A C2-aperture of 100 μm was selected for the PB41 data collection. Coma free alignment was performed with AutoCtf/Sherpa. Data were acquired in parallel illumination mode using EPU (Thermo Fisher Scientific) software at a nominal magnification of 130kx (1.041 Å pixel size). Both datasets for the HAdV-F41 structure at pH=7.4 were collected in super-resolution mode. Due to a preferred orientation of PB41, a second data set was collected at a 30° tilted stage. Data collection parameters are listed in the Supplementary Table 7.

### Data processing and structure determination HAdV-F41 pH=7.4

Two datasets were collected on HAdV-F41 pH=7.4 and initially processed independently. Data were initially processed using Relion 3.0(50) and continued in Relion 3.1 (beta)(51). Beam-induced motion was corrected using Relion’s MotionCor2(52) implementation, at which step the super-resolution movies were binned once, and the per-micrograph CTF estimated using GCTF(53) for all data sets. Particles were manually picked and subjected to reference-free 2D classification and well-resolved classes were combined and subjected to 3D classification, applying icosahedral symmetry (I3 according to Crowther(54)) and a mask of the capsid structures. A low-pass filtered (50 Å) volume of HAdVC-5 (EMD-7034(20)) was used as a reference volume. Particles were classified into two classes, resulting in 99% of particles allocated to one well-resolved class which was used for downstream processing. 3D refinement was performed using the output of the 3D classification as a reference model, low-pass filtered to 50 Å, with no additional Fourier-padding. Following refinement, data were post-processed, and the particles subjected to per-particle CTF refinement, Bayesian polishing and another round of per-particle CTF refinement. The particles were subjected to an additional round of 3D refinement before combining both datasets and performing a final 3D refinement, with no additional Fourier-padding. The resolution was calculated using the gold standard FSC (threshold 0.143) to 3.84 Å after postprocessing. Finally, the data were corrected for the Ewald’s sphere curvature using Relion, which led to a local improvement of the electron density map with and a new average resolution of 3.77 Å. Local Resolution estimates were calculated using ResMap(55).

A homology model was generated using the SWISS-MODEL server(56) the HAdV-F41 capsid protein sequences, for which homologues have been structurally determined. The resulting homology model was based on the reported HAdV-D26 structure (PDB: 5TX1(21)). The model was manually docked into the HAdV-F41 electron density in ChimeraX(57) and the map corresponding to the asymmetric unit (ASU) extracted. The ASU map was locally sharpened using Phenix’s(55) autosharpen tool. Subsequently, the HAdV-F41 homology model was docked and subjected to an initial round of real-space-refinement using Phenix. The structure was fully refined using iterative cycles of Phenix’s real-space-refinement and model building in Coot(59).

### Localized asymmetric reconstruction

To improve the map quality surrounding the penton base monomer and the HVR2-loop containing region, the map was improved using the localized asymmetric reconstruction workflow reported by Ilca *et al*(37) and implemented in Scipion v2.0(60). Coordinates for the sub-particles were determined in ChimeraX and subsequently located by applying icosahedral symmetry and extracted in Scipion v2.0. Sub-particles were subsequently filtered to exclude particles not present within a [-20°, 20°] range from the image plane. The resulting sub-particles were then subjected to an asymmetric 3D classification. To increase the probability of convergence during classification, changes in the origins and orientations were not allowed. A subsequent 3D refinement yielded a 3D reconstruction of the penton base monomer and the HVR2-loop containing region to a resolution of 3.0 Å and 3.35 Å, respectively. Average resolutions were calculated according to the gold standard FSC calculations (threshold 0.143). Data processing statistics for the asymmetric localized reconstruction are given in Supplementary Table 8. 3DFSC curves were calculated using the Remote 3DFSC processing server(61).

### Image processing and model building for HAdV-F41 at pH=4.0

Data were processed as described for the HAdVF-41 pH=7.4 structure up until the first 3D refinement. The volume HAdVF-41 at pH=7.4 was low-passed filtered to 10 Å and used as a reference. The resolution was estimated to 5.0Å using the gold standard FSC (threshold 0.143) after postprocessing. Local Resolution estimates were calculated using ResMap.

The HAdV-F41 pH=7.4 model was fitted into the reconstructed HAdV-F41 pH=4.0 density using ChimeraX and an ASU extracted and the resulting map locally sharpened using Phenix. The model was then further fitted and energy minimized using Namdinator(62).

### Data processing and structure determination recombinant HAdV41 penton base

The HAdV-F41 penton base (PB41) data (untilted and tilted at 30°) were processed using Relion 3.1 (beta), with beam-induced motion correction and CTF estimation performed as for the HAdV-F41 structure. Particles were picked using the automated particle picker crYOLO(63) using the available Phosaurus generalized model. Reference-free 2D classification of particles was performed in Relion and revealed a significant proportion of particles with the same view, suggesting a preferred orientation of the specimen. From the 0° data, an initial model was generated in cryoSPARC(64). Well-resolved 2D classes were combined and subjected to 3D refinement using the same reference model as during 3D classification, low pass filtered to 10 Å. Following refinement, data were post-processed, and the particles subjected to per-particle CTF refinement, Bayesian polishing and another round of per-particle CTF refinement before performing a final round of 3D refinement. Inspection of the final volume revealed poor resolution along one of the axes (Supplementary Figure 6). We therefore collected data on a tilted specimen stage. As data collection at a tilted stage leads to a defocus gradient along the image path, per-particle CTF refinement was performed after particle extraction and before reference-free 2D classification, using GCTF. Subsequent processing steps were performed as described for the data collected on an untilted specimen stage. The average resolutions were estimated to 3.4 Å (untilted) and 3.7 Å (30° tilt), using the gold standard FSC (threshold 0.143) after postprocessing. Local Resolution estimates were calculated using ResMap. 3DFSC curves were calculated using the Remote 3DFSC processing server(61).

The PB41 volume generated from the data collected on the tilted stage was used for down-stream model building and model refinement. The penton base monomer chain from the HAdV-F41 pH=7.4 model was initially fitted into PB41 volume using Namdinator(62) and outlying residues pruned in Coot. Subsequently the model was fully built and refined using iterative cycles of real-space-refinement in Phenix and model building in Coot.

### Calculation of surface charge distribution

Surface charges for HAdV-C5 (PDB: 6CGV(19)), HAdV-D26 (PDB: 5TX1(21)) and HAdV-F41 was calculated using the pdb2pqr-apbs software package(65) at pH=7.4 and pH=4.0.

Bioinformatics workflow to determine the protein identity of the unknown chain For each direction of the unknown chain, a 24mer poly-alanine chain was manually placed into the respective density and initially real-space-refined using Coot. A list of sequences was screened by using a job pipeline including a mutation step in Coot and real-space-refinement in Phenix. Custom bash scripts written for this purpose are available upon request. An extended description is given in Supplementary Methods 1.

### Molecular graphics and visualization

Figures of protein structures and electron densities were generated using ChimeraX.

## Acknowledgments

**General:** We are grateful to Dr. Paul Emsley for help with Coot scripting, Dr. Michael Hall for help with cryoSPARC, and Tom Terwilliger for help with Phenix. We thank Stefan Nord, Jonas Naslund and Andreas Sjodin (FOI, Swedish Defence Research Institute, Umea, Sweden) for help with DNA sequencing. Cryo-EM data were collected at the Umea Core Facility for Electron Microscopy (SciLifeLab National Cryo-EM facility and part of National Microscopy Infrastructure, NMI VR-RFI 2016-00968), supported by instrumentation grants from the Knut and Alice Wallenberg Foundation and the Kempe Foundations. Protein identification was performed at the Proteomics Core Facility of Sahlgrenska Academy, University of Gothenburg. We thank the High Performance Computing Center North (HPC2N) at Umea University for providing computational resources and valuable support during test and performance runs (SNIC projects 2018/5-185, 2019/3-400,-668,5-76 & 5-158).

## Author contributions

K.R., A.L., N.A. and L.-A.C. conceived and designed the study. A.L. produced and purified virus particles. A.R. purified recombinant penton base protein. K.R. collected cryo-EM data, performed image processing, model building and validation. K.R., A.L., N.A. and L.-A.C. interpreted structural data. J.F. performed proteomics analysis. K.R., A.L., N.A. and L.-A.C wrote the original manuscript with input from J.F. All authors reviewed and approved the final manuscript.

## Data and materials availability

The scripts used for the bioinformatics analysis are available upon request. Coordinates reported in this study have been deposited with the Protein Data Bank with accession codes XXXX (HAdV-F41 asymmetric unit) and XXXX (HAdVF-41 (free) penton base). Electron microscopy maps and half-maps have been deposited in the Electron Microscopy Data Bank with the accession codes EMD-YYYYY (HAdV-F41 pH=7.4), EMD-YYYYY (HAdV-F41 pH=4.0), EMD-YYYYY (HAdV-F41 (free) penton base), EMD-YYYYY (localized asymmetric reconstruction of the HAdV-F41 penton base), EMD-YYYYY (localized asymmetric reconstruction of the HAdV-F41 HVR2-containing loop).

