## Supplementary Information for "The structure of enteric human adenovirus 41 - a leading cause of diarrhea in children"

### Supplementary Materials

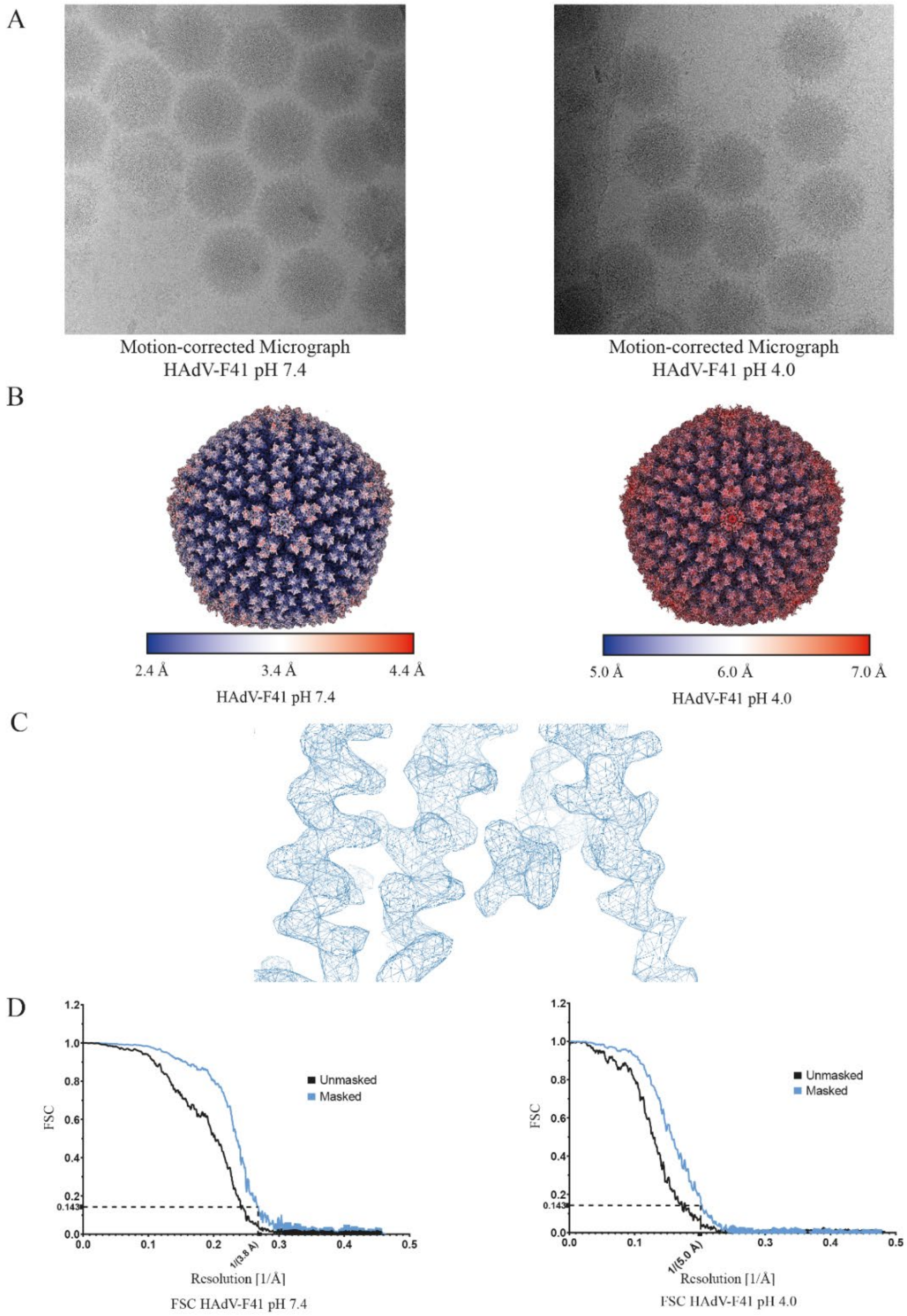


**Supplementary Figure 1: HAdV-F41 structures at pH=7.4 and pH=4.0.** (A) Motion-corrected micrograph of HAdV-F41 particles at pH=7.4 and pH=4.0. (B) Surface representation of HAdV-F41 at pH=7.4 and pH=4.0. Colouring represents local resolution of the structure. (C) Close up of the HAdV-F41 pH=7.4 density, shown as a blue mesh. (D) Graphs showing the Fourier-Shell-Correlation (FSC) vs. resolution [1/Å] of the unmasked and masked volumes. FSC=0.143 is indicated as a black dashed line.

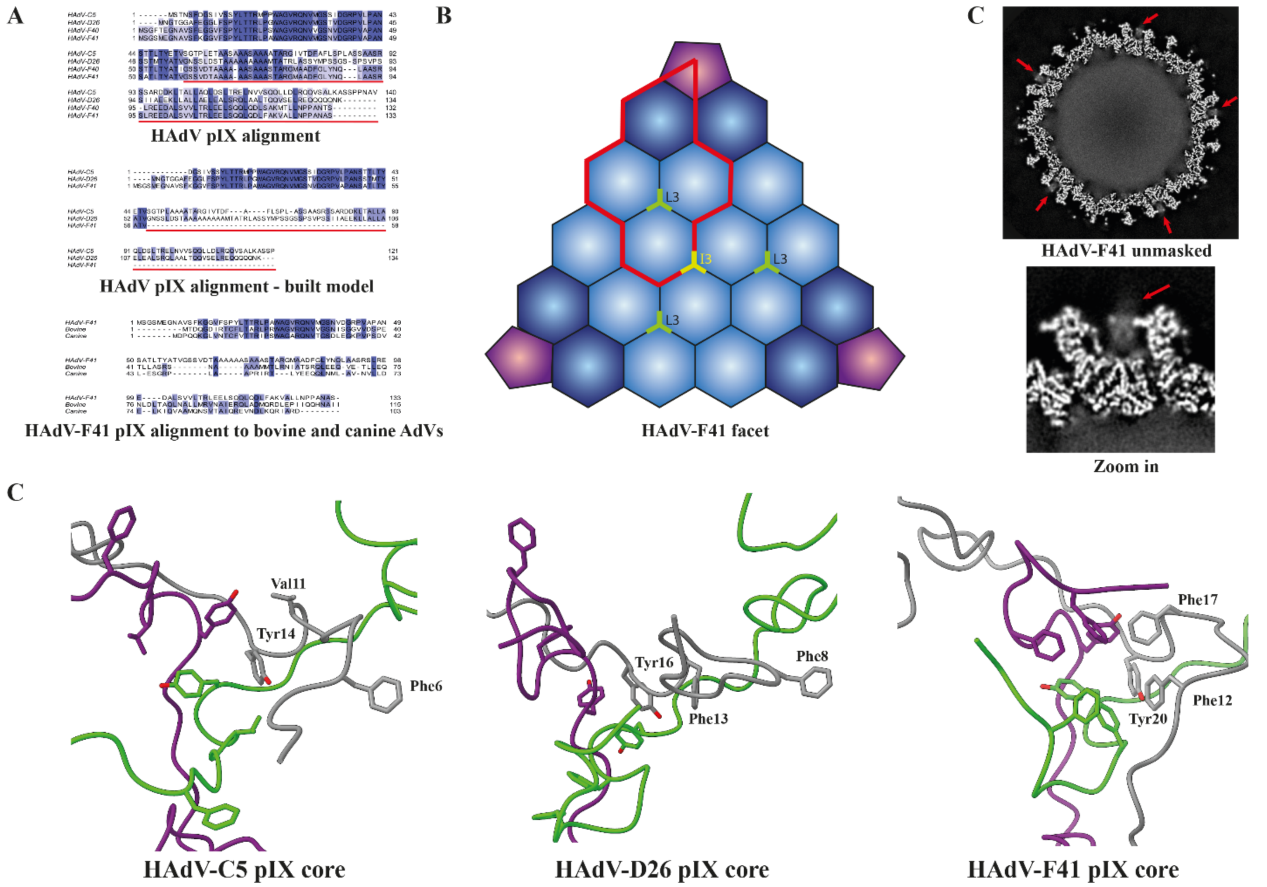


**Supplementary Figure 2: Protein IX.** (A) *Top* Sequence alignments of HAdV-C5, -D26, -F40 and –F41 pIX. *Centre* Sequence alignment of pIX chains built in the structures of HAdV-C5 (PDB: 6B1T(*20*)), HAdV-D26 (PDB: 5TX1(*21*)) and HAdV-F41. *Bottom* Sequence alignment of HAdV-F41 pIX with bovine and canine AdVs. Colouring for all alignments represents % sequence identity with dark blue representing 100% homology. The red bar represents the C-terminus of pIX (pIX-C). (B) Schematic representation of one facet of the adenovirus capsid. Penton bases, peripentonal hexons and hexons are coloured as in Fig. 1. The thick red line represents the asymmetric unit of the capsid. The pIX-N triskelia located at the local 3-fold symmetry axis ("L3") and the icosahedral symmetry axis ("I3") are coloured green and yellow, respectively. (C) Computational slice through the unmasked HAdV-F41 volume and a zoom in on the non-peripentonal hexon space. The delocalised density pointed out with an arrow is proposed to represent pIX-C. (D) Graphical representation of the hydrophobic core of pIX of HAdV-C5, -D26 and –F41. The three chains are shown as cartoon, with the conserved hydrophobic core residues in HAdV-F41 shown in stick representation.

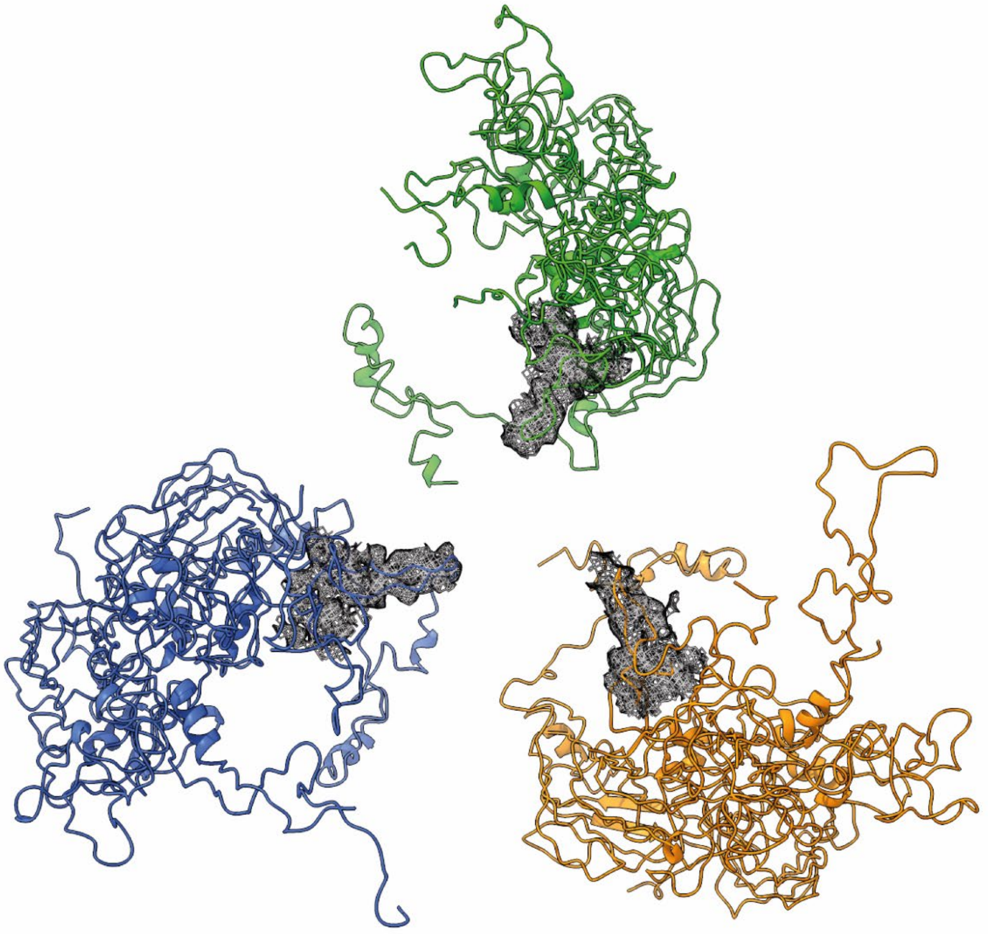

**Supplementary Figure 3: Non-peripentonal hexon HVR2s point towards a pseudo-threefold symmetry axis.** The pseudo-trifold symmetry created by three non-peripentonal hexon chains with the HVR2-containing loops pointing towards the center of the symmetry. The hexon chains are shown in cartoon representation and the electron density corresponding to the HVR2-containing loops is shown as as a black mesh.

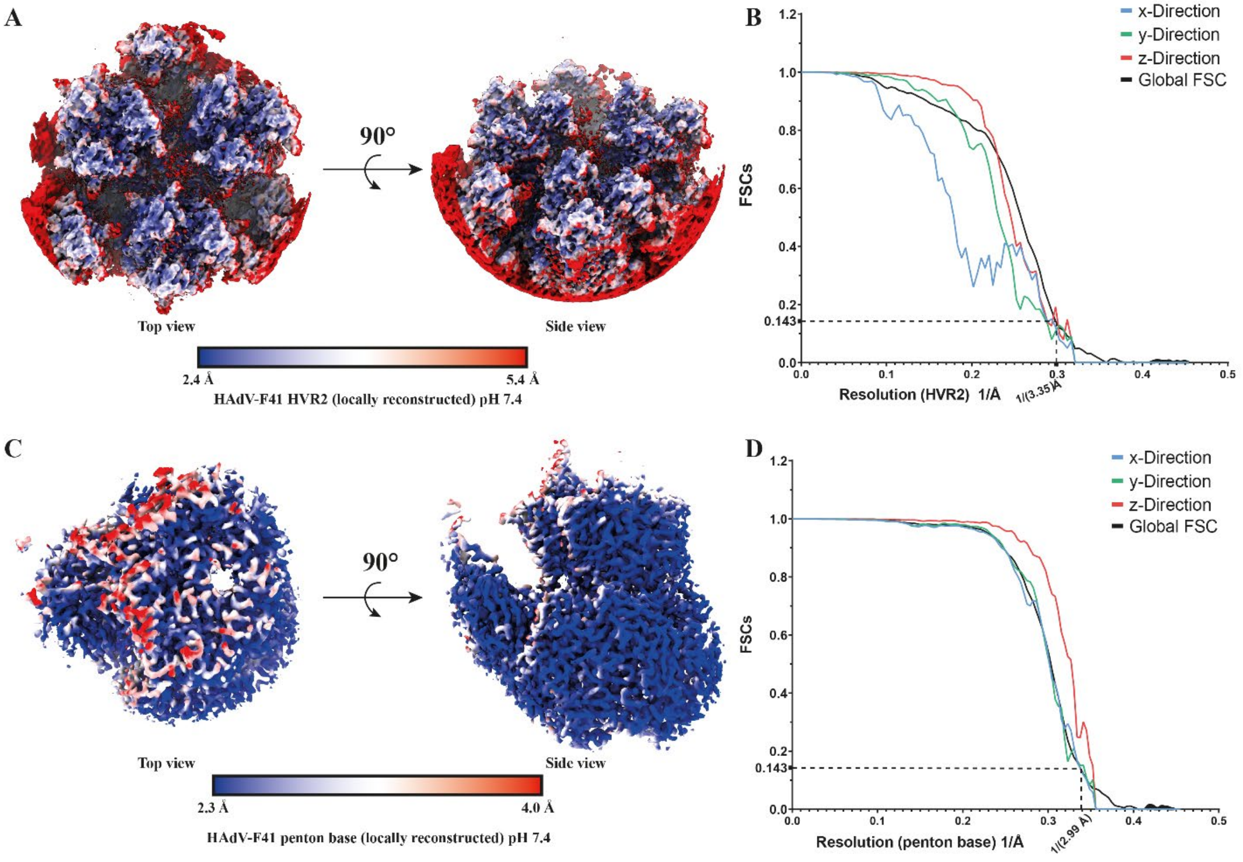


**Supplementary Figure 4: Localised asymmetric reconstructions of HAdV-F41 features**. (A) Locally reconstructed volumes of the HVR2-loop containing region coloured by local resolution estimates. (B) 3D Fourier-Shell-Correlation (3DFSC) curves of the locally reconstructed volume. The average resolution at FSC(global)=0.143 is indicated by a black dashed line. (C) Locally reconstructed volume of the penton base coloured by local resolution estimates. (D) 3D Fourier-Shell-Correlation (3DFSC) curves of the locally reconstructed volume. FSC=0.143 is indicated by a black dashed line.

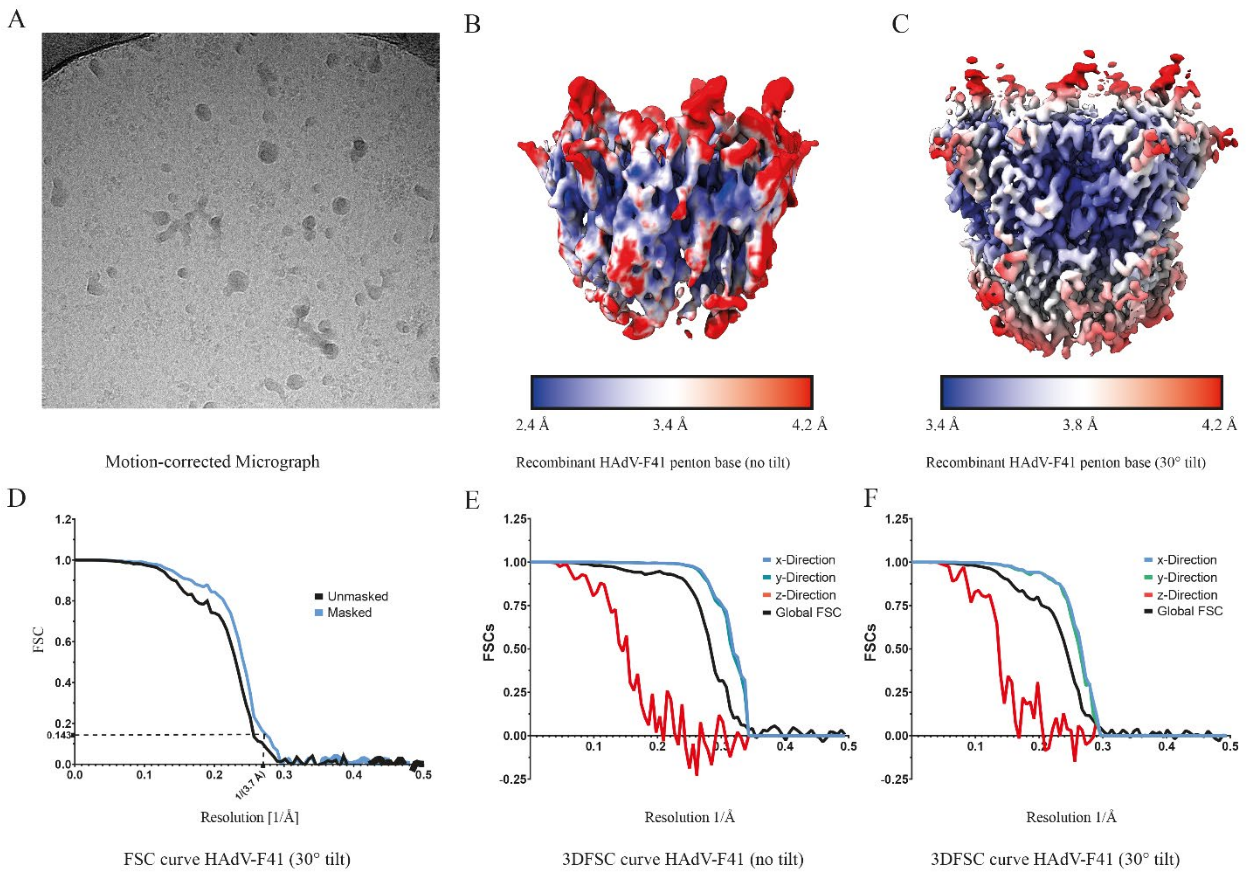


**Supplementary Figure 5: 3D reconstruction of the free penton base homopentamer at 0º and 30º stage tilt.** (A) Motion-corrected micrograph of recombinant HAdV-F41 penton base. (B) Surface representation of the recombinantely expressed penton base collected at no tilt of the specimen stage. Colouring represents local resolution estimations. (C) Surface representation of the recombinantely expressed penton base collected at 30° tilt of the specimen stage. Colouring represents local resolution estimations. (D) FSC curves of the unmasked and masked volumes of the HAdV-F41 penton base collected at a 30° tilt. The average resolution at FSC=0.143 is indicated by a black dashed line. (E) 3D Fourier shell correlation (3DFSC) curves of the locally reconstructed volumes collected at no tilt of the specimen stage. (F) 3D Fourier shell correlation (3DFSC) curves of the locally reconstructed volumes collected at 30° tilt of the specimen stage.

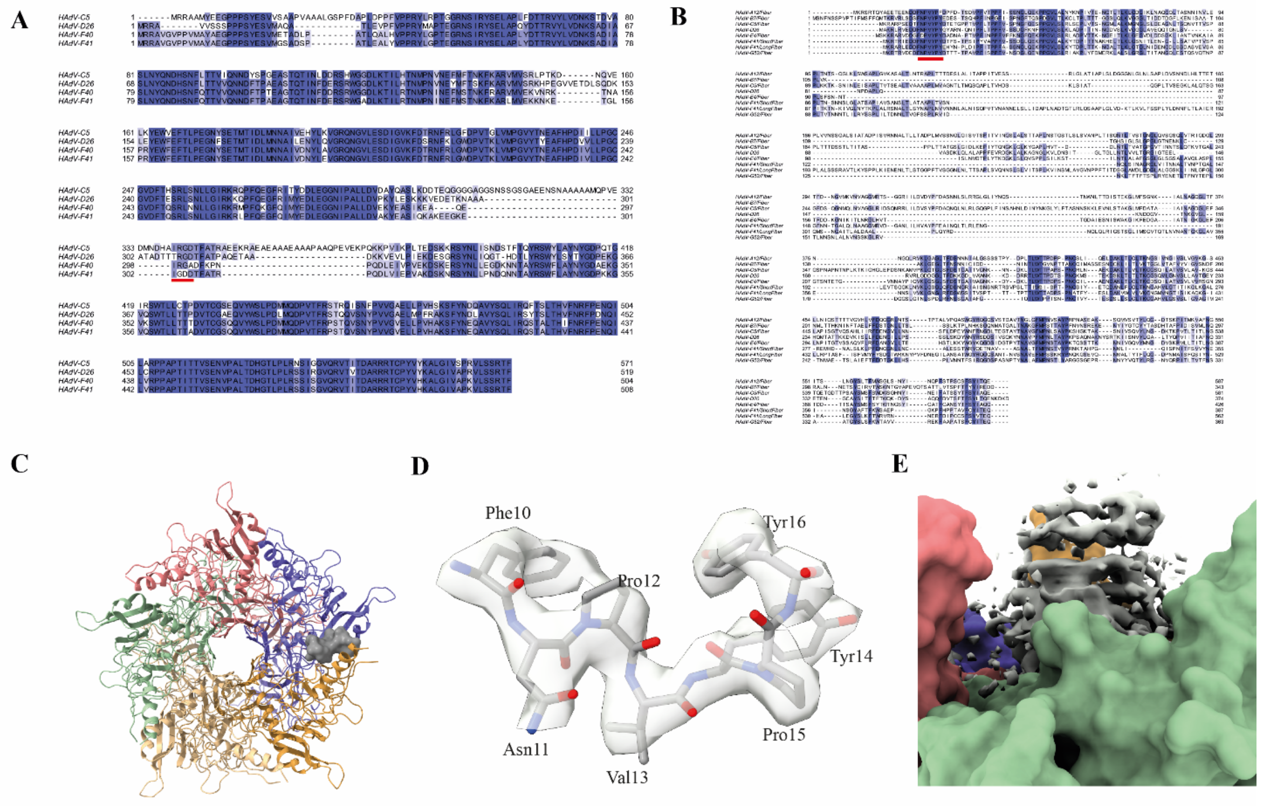


**Supplementary Figure 6: The virion-bound penton base and its interface with the fibre tail.** (A) Alignment of the HAdV-C5, -D26, -F40 and –F41 penton base amino acids sequences. Colouring reflects % sequence identity. The red bar represents the RGD/IGDD motif. (B) Alignment of the fibre tail sequences from a representative of each HAdV species (A-G). Colouring reflects % sequence identity. The red bar represents the conserved fibre tail motif. (C) Cartoon representation of the HAdV-F41 virion-bound penton base in complex with the HAdV-F41 fibre tail. Individual penton base chains are shown as cartoons and coloured in green, yellow, orange, pink and purple. One of the fibre tail electron densities is shown as a grey surface. (D) The built fibre tail model and the corresponding electron density. The fibre tail is shown as grey sticks and the electron density is shown as semi-transparent surface. (E) HAdV-F41 fibre density. The density is shown as a grey surface and the virus-bound penton base as coloured surfaces.

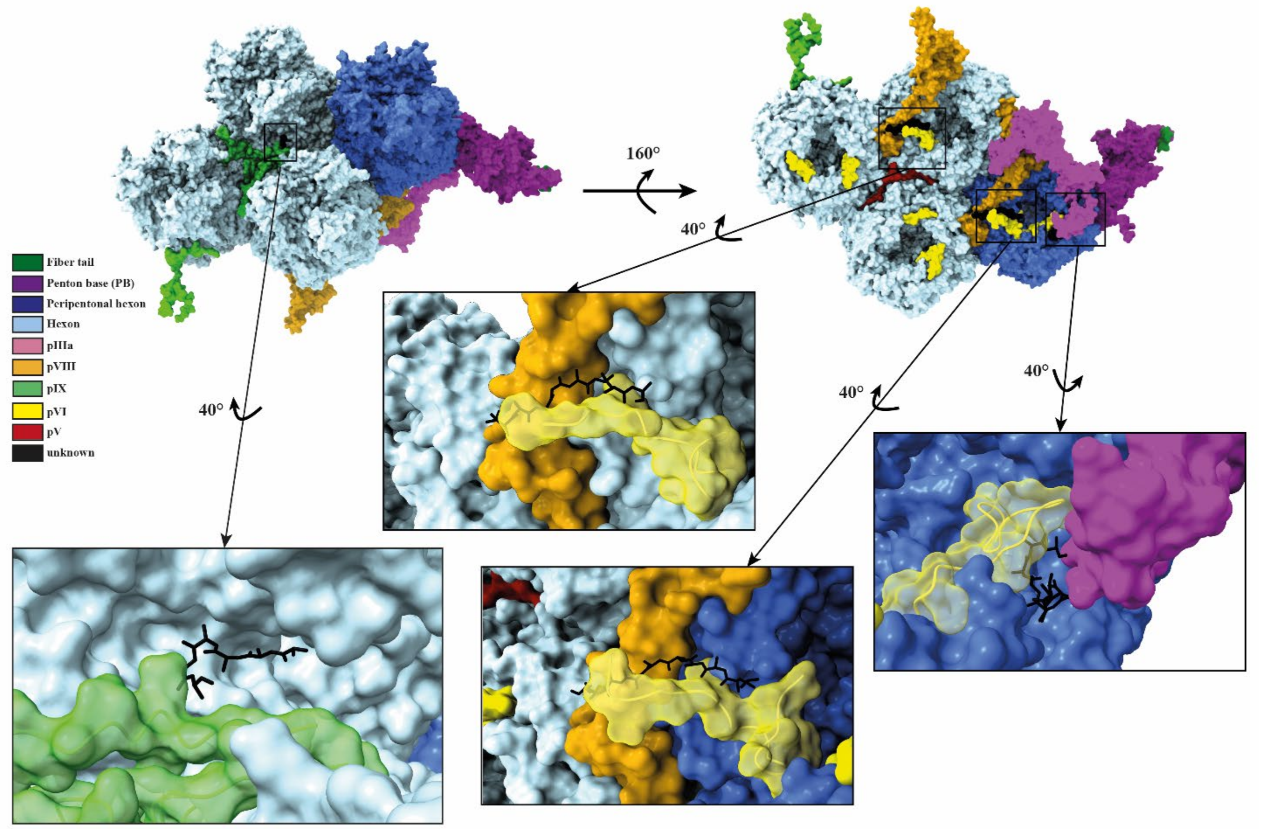


**Supplementary Figure 7: Location and interactors of four unidentified peptide chains.** Surface representation of the HAdV-F41 asymmetric unit coloured according to the given legend. A close-up of each individual unknown chain (black) is shown as individual panels. In each the unidentified peptide chain is shown as black sticks covered by a semi-transparent surface. The surrounding interacting proteins are shown as cartoons covered by a semi-transparent surface.

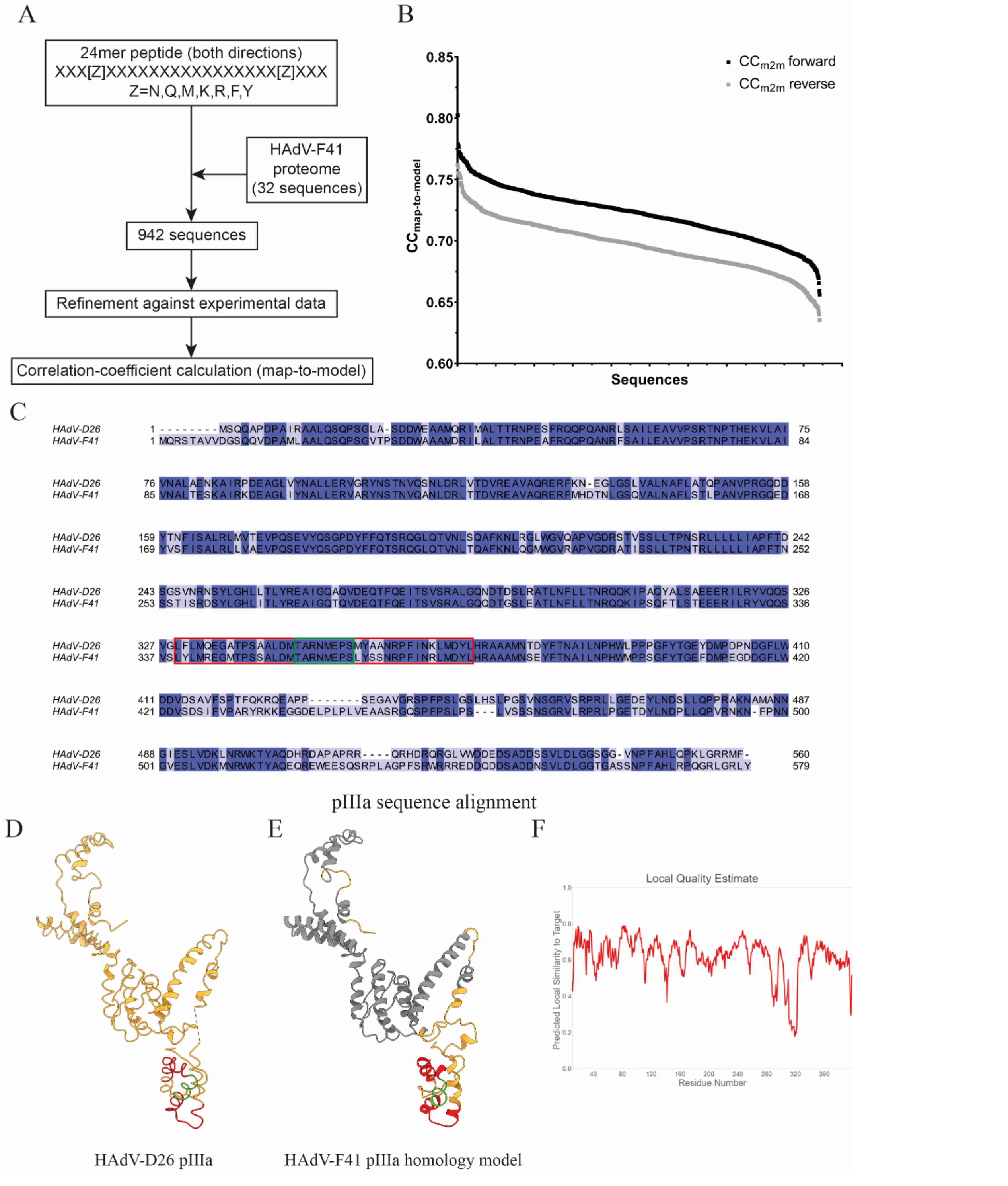


**Supplementary Figure 8: Bioinformatics workflow to determine identity of the unknown chain.** (A) Flowchart of the bioinformatics workflow steps. (B) Graph showing the $CC_{map\text{-}to\text{-}model}$ distribution for the sequences screened using the bioinformatics workflow. (C) Sequence alignment of HAdV-D26 and HAdV-F41 pIIIa. Identical residues are shown on a dark blue background. The two identified sequence candidates are marked with a red box and the overlapping sequence with a green box. (D) Cartoon representation (gold) of HAdV-D26 pIIIa (PDB: 5TX1(*21*)). The sequences marked with boxes in C are indicated the model with the same colours as those boxes. (E) The homology model generated for HAdV-F41 by the SWISS-MODEL server(*56*). The colouring represents the part of the protein that could be modelled in the HAdV-F41 structure (grey), the model for which no density was visible in the HAdV-F41 structure (gold) and the pIIIa sequences identified as potential candidates are shown in red with the overlapping residues coloured in green. (F) Plot showing the per-residue model quality generated by SWISS-MODEL server.

**Supplementary Table 1:** List of validated HAdV-F41 proteins detected by mass spectrometry after tryptic and chymotryptic digest.

| | Description | Coverage [%] | # Peptides | # Peptide Spectrum Matches | MW [kDa] |
|---|---|---|---|---|---|
| trypsin | 100k | 33.98 | 21 | 29 | 87.3 |
| | E1B 19k | 10.00 | 2 | 3 | 19.6 |
| | 52-55k | 50.66 | 20 | 73 | 42.9 |
| | E1B 55k | 8.47 | 4 | 5 | 52.1 |
| | E2A DBP | 50.63 | 21 | 44 | 53.6 |
| | E2B preterminal protein | 44.48 | 24 | 69 | 72.0 |
| | E3 14.7k | 25.41 | 3 | 4 | 14.0 |
| | E4 hypothetical protein 2 | 13.85 | 1 | 1 | 14.9 |
| | hexon | 69.95 | 61 | 2336 | 103.9 |
| | IIIa | 70.29 | 32 | 442 | 64.7 |
| | IVa2 | 62.56 | 23 | 89 | 50.9 |
| | IX | 100.00 | 10 | 204 | 13.6 |
| | L2 pMu | 13.04 | 2 | 9 | 7.6 |
| | long fiber protein | 45.37 | 16 | 54 | 60.6 |
| | penton base | 67.91 | 24 | 241 | 57.0 |
| | pVII | 42.08 | 11 | 473 | 20.3 |
| | pVIII | 54.94 | 8 | 80 | 25.3 |
| | short fiber protein | 46.25 | 11 | 55 | 41.4 |
| | truncated U exon protein | 62.26 | 3 | 6 | 6.4 |
| | V | 78.16 | 31 | 353 | 39.7 |
| | VI | 66.92 | 16 | 487 | 29.1 |
| | **Description** | **Coverage [%]** | **# Peptides** | **# Peptide Spectrum Matches** | **MW [kDa]** |
| chymotrypsin | 100k | 8.11 | 4 | 4 | 87.3 |
| | 52-55k | 42.22 | 19 | 34 | 42.9 |
| | E2A DBP | 19.62 | 9 | 13 | 53.6 |
| | E2B preterminal protein | 17.44 | 12 | 16 | 72.0 |
| | E3 14.7k | 6.56 | 1 | 1 | 14.0 |
| | hexon | 84.00 | 151 | 1075 | 103.9 |
| | IIIa | 75.99 | 70 | 227 | 64.7 |
| | IVa2 | 32.06 | 15 | 24 | 50.9 |
| | IX | 76.69 | 13 | 44 | 13.6 |
| | L2 pMu | 10.14 | 1 | 3 | 7.6 |

| | | | | | |
|---|---|---|---|---|---|
| | long fiber protein | 68.86 | 51 | 85 | 60.6 |
| | penton base | 71.65 | 42 | 141 | 57.0 |
| | pVII | 11.48 | 1 | 27 | 20.3 |
| | pVIII | 34.76 | 9 | 19 | 25.3 |
| | short fiber protein | 68.22 | 34 | 58 | 41.4 |
| | V | 23.56 | 8 | 27 | 39.7 |
| | VI | 52.26 | 15 | 50 | 29.1 |

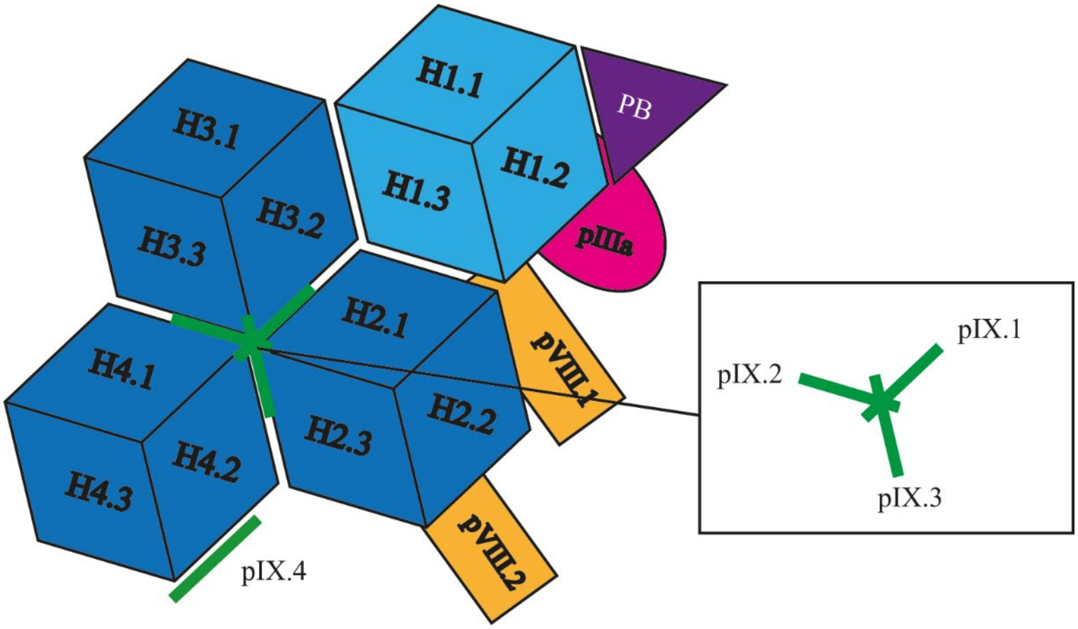


**Ass. Figure 1:** Assignment of Capsid protein identifiers listed in Supplementary Table 2.

**Supplementary Table 2:** PISA(*66*) analysis of HAdV-F41 capsid protein interactions. Protein assignment is shown in associate Figure S1. Identity of minor capsid protein pIVs and unknown chains are assigned based on their chain ID in the deposited structure. Area = interacting surface area, ΔG = Gibbs free-energy, Nhb = number of hydrogen bonds, Nsb = number of salt bridges.

| Capsid protein 1 | Capsid protein 2 | Area [$Å^2$] | □G | Nhb | Nsb |
|---|---|---|---|---|---|
| H4.3 | H4.1 | 8316.6 | -92.7 | 71 | 7 |
| H4.3 | H4.2 | 8044.1 | -89.5 | 80 | 12 |
| H3.2 | H3.1 | 7996.9 | -79.2 | 89 | 10 |
| H2.2 | H2.1 | 7942.2 | -91 | 73 | 10 |
| H4.2 | H4.1 | 7650.4 | -79.5 | 79 | 5 |
| H3.3 | H3.1 | 7515.2 | -86.4 | 71 | 11 |
| H3.3 | H3.2 | 7413.8 | -84.6 | 59 | 10 |
| H2.3 | H2.1 | 7269.5 | -87 | 54 | 11 |
| H1.3 | H1.1 | 6926.2 | -79.1 | 60 | 11 |
| H2.3 | H2.2 | 6907.9 | -77.9 | 67 | 5 |
| H1.3 | H1.2 | 6576.9 | -74.4 | 57 | 5 |
| H1.2 | H1.1 | 5942.4 | -62.6 | 48 | 8 |
| H2.2 | H3.1 | 4148.2 | -18.6 | 118 | 24 |
| pIIIa | H3.1 | 2817.7 | -13.6 | 73 | 4 |
| H2.2 | H3.3 | 1948.6 | -8.7 | 63 | 10 |
| pVIII.2 | H4.1 | 1672.1 | -6.2 | 33 | 0 |
| pVIII.1 | H3.1 | 1494.1 | -12.9 | 24 | 6 |
| H2.3 | H3.3 | 1425.9 | -6.4 | 22 | 2 |
| pVIII.1 | H1.3 | 1393 | -9.5 | 11 | 3 |
| pVIII.2 | H2.3 | 1373.3 | -7.4 | 14 | 5 |
| Penton base | H1.2 | 1314.8 | -2.5 | 11 | 2 |
| pVIII.1 | H2.2 | 1233.2 | -14.5 | 19 | 2 |
| pIX.3 | H4.2 | 1232.3 | -11.1 | 11 | 3 |
| pIX.2 | H3.3 | 1210.4 | -9.7 | 12 | 1 |

| | | | | | |
|---|---|---|---|---|---|
| pIX.3 | H2.1 | 1095.2 | -9.6 | 10 | 0 |
| pIX.2 | H4.2 | 879.1 | -8.9 | 9 | 0 |
| pIX.1 | H2.1 | 1059.4 | -8.9 | 7 | 3 |
| pVIII.1 | pIIIa | 1007.2 | -11.4 | 4 | 0 |
| H2.1 | H4.2 | 910.8 | -7.7 | 7 | 0 |
| H2.1 | H3.3 | 850.8 | -6.8 | 5 | 1 |
| H3.2 | H1.1 | 842.3 | -2.9 | 5 | 2 |
| H4.2 | H3.3 | 806 | -1.4 | 14 | 1 |
| pV | H3.3 | 907 | -5.2 | 10 | 2 |
| pIIIa | H1.2 | 892.6 | -5.3 | 10 | 2 |
| pVIII.1 | H1.2 | 869.9 | -10.4 | 10 | 2 |
| pVIII.2 | H2.2 | 842.5 | -9.9 | 9 | 0 |
| pV | H2.1 | 800.8 | -10.2 | 7 | 0 |
| pVIII.2 | H4.2 | 736.9 | -3.9 | 7 | 4 |
| pVIII.1 | H3.2 | 622.7 | -6.8 | 2 | 0 |
| pIX.1 | pIX.3 | 703.9 | -11.2 | 3 | 0 |
| H1.3 | pVI | 657.5 | -4.8 | 7 | 2 |
| H2.2 | H1.3 | 601.2 | -4.8 | 2 | 1 |
| H3.2 | H1.3 | 600.4 | -1.2 | 2 | 0 |
| H2.3 | H4.2 | 597.3 | -1.8 | 0 | 4 |
| H4.1 | H3.3 | 589.1 | -1.5 | 1 | 2 |
| H2.1 | H3.2 | 570 | 1.1 | 1 | 3 |
| H1.1 | pVI | 584.2 | -3.5 | 5 | 0 |
| pIIIa | H1.1 | 554.5 | -7.5 | 1 | 0 |
| H2.1 | H1.3 | 520.3 | -1.3 | 4 | 2 |
| pIX.2 | pIX.3 | 487.1 | -9.7 | 2 | 0 |
| H1.2 | pVI | 427.2 | -3.6 | 2 | 1 |
| H4.3 | pVI | 423 | 0 | 3 | 0 |
| pV | H4.2 | 404.7 | -1.5 | 1 | 0 |
| pIX.1 | pIX.2 | 392.8 | -6.8 | 2 | 0 |
| H2.1 | H3.1 | 380.7 | -5.2 | 2 | 0 |
| H4.1 | pVI | 370.2 | -3.4 | 1 | 2 |
| H4.2 | pVI | 368.1 | -1.8 | 2 | 2 |
| H2.3 | pVI | 365.3 | -5.4 | 3 | 0 |
| H3.1 | pVI | 362.4 | -2.4 | 7 | 0 |
| H2.1 | pVI | 359.6 | -3.2 | 2 | 2 |
| H4.3 | pIX.4 | 357.9 | -4.2 | 4 | 0 |
| H1.3 | pVI | 354.6 | -2.2 | 4 | 0 |
| pVIII.1 | H2.1 | 353.6 | -5.6 | 4 | 0 |
| H2.3 | H3.1 | 353.5 | -0.1 | 7 | 0 |
| H3.3 | pVI | 343.8 | -1.9 | 2 | 0 |
| H4.1 | pVI | 343 | -3.4 | 2 | 2 |
| H2.2 | H3.2 | 341.6 | -2.6 | 3 | 0 |
| pIX.1 | H3.3 | 335.1 | -2.9 | 2 | 0 |
| pIX.3 | H3.3 | 333.5 | -4.7 | 3 | 0 |
| pIX.2 | H2.1 | 242.4 | -3.7 | 2 | 0 |

| | | | | | |
|---|---|---|---|---|---|
| pV | H3.2 | 328.9 | -2.6 | 2 | 1 |
| Unknown Y | H3.3 | 326.2 | -3.1 | 3 | 0 |
| H1.2 | pVI | 310.8 | -4.1 | 2 | 0 |
| Fibre tail | Penton base | 265.9 | -4.7 | 1 | 0 |
| H3.2 | pVI | 263.1 | -2.4 | 1 | 0 |
| H2.2 | pVI | 262.5 | -1.8 | 3 | 0 |
| Unknown V | pVIII.2 | 242.1 | -2.5 | 6 | 0 |
| H4.3 | pVI | 236 | -2.5 | 0 | 0 |
| Unknown X | pVIII.1 | 229 | -3.3 | 4 | 0 |
| H4.2 | pVI | 228.9 | -3.1 | 1 | 0 |
| Unknown X | pVI | 226.8 | -3.2 | 3 | 0 |
| Unknown V | pVI | 225.4 | -3.8 | 4 | 0 |
| pIX.1 | H4.2 | 222.5 | -3.3 | 2 | 0 |
| Unknown W | pVI | 221.8 | -3.1 | 2 | 0 |
| H3.1 | pVI | 215.6 | -2.8 | 0 | 0 |
| Unknown W | H1.3 | 184.5 | -2.2 | 0 | 0 |
| Unknown W | H1.1 | 149.9 | -2.4 | 0 | 0 |
| pVIII.2 | H2.1 | 145.2 | 0 | 0 | 0 |
| pVIII.1 | H1.1 | 105.5 | -0.4 | 0 | 0 |
| Penton base | H1.1 | 143.9 | -2.1 | 0 | 0 |
| pV | H2.3 | 142.7 | -4.8 | 0 | 0 |
| pVIII.2 | H4.1 | 136.6 | -2.1 | 1 | 0 |
| pVIII.1 | H3.1 | 108.9 | -2.3 | 2 | 0 |
| Unknown W | pIIIa | 136 | -1 | 1 | 0 |
| pIIIa | H2.1 | 112.1 | -1.4 | 1 | 0 |
| pVIII.2 | H3.3 | 106.8 | 0 | 1 | 0 |
| pIX.2 | H4.1 | 87.1 | -1.9 | 0 | 0 |
| pIX.3 | H2.3 | 58.3 | -1.4 | 0 | 0 |
| pV | H4.1 | 86.4 | -1.2 | 0 | 0 |
| pIX.3IIIa | H3.2 | 82.1 | 0.9 | 0 | 0 |
| Unknown X | H1.2 | 80.3 | 0.6 | 2 | 0 |
| pIX.3IIIa | H2.2 | 76.7 | -1.1 | 0 | 0 |
| pIX.3IIIa | Penton base | 71.9 | -1.4 | 0 | 0 |
| Unknown Y | pIX.3 | 68.7 | -1.1 | 0 | 0 |
| Unknown V | H2.2 | 65.4 | -0.7 | 0 | 0 |
| pIIIa | pVI | 59.5 | 0.4 | 1 | 0 |
| pIIIa | H3.3 | 58.6 | 0 | 0 | 0 |
| pVIII.2 | pVI | 52.9 | -1.7 | 0 | 0 |
| H2.2 | H1.2 | 49.5 | 0.6 | 2 | 0 |
| pVIII.1 | pVI | 47.1 | -0.4 | 0 | 0 |
| H2.1 | pVI | 42 | 0.3 | 1 | 0 |
| H4.1 | pVI | 37.4 | 0.8 | 0 | 0 |
| Unknown W | H1.2 | 36.4 | -0.5 | 0 | 0 |
| H1.1 | H4.2 | 35.5 | 1.4 | 1 | 0 |
| H2.3 | pVI | 33.9 | -0.2 | 0 | 0 |
| H2.3 | H3.2 | 29.3 | -0.4 | 0 | 0 |

| | | | | | |
|---|---|---|---|---|---|
| H4.1 | pVI | 29.2 | -0.4 | 0 | 0 |
| Unknown Y | H3.2 | 27.4 | -0.6 | 0 | 0 |
| H1.1 | pVI | 27.2 | -0.4 | 0 | 0 |
| pVI | pVI | 26.6 | 0.5 | 0 | 0 |
| H4.3 | pVI | 26.5 | 0.1 | 0 | 0 |
| pVIII.2 | H3.1 | 24.2 | 0.1 | 1 | 0 |
| H4.2 | pVI | 20.7 | -0.1 | 0 | 0 |
| H3.2 | pVI | 20.6 | -0.1 | 0 | 0 |
| H3.3 | pVI | 18.1 | 0 | 0 | 0 |
| H4.2 | H3.2 | 5.9 | 0 | 0 | 0 |

**Supplementary Table 3:** Protein sequence identity between HAdV-F41 and HAdV-C5 and HAdV-D26 as well as the overall root mean square deviation (RMSD) after structure alignment.

| | HAdV-C5 [%] | RMSD [Å] (PDBID 6B1T) | HAdV-D26 [%] | RMSD [Å] (PDBID 5TX1) |
|---|---|---|---|---|
| **Hexon** | 77.5 | 2.67 | 78.9 | 0.96 |
| **Penton Base** | 68.8 | 1.8 | 76.8 | 2.88 |
| **Fibre short** | 38.1 | N/A | 30.9 | 0.6 |
| **Fibre long** | 32.6 | N/A | 42.7 | 0.6 |
| **pIIIa** | 70.1 | 2.2 | 74.1 | 1.67 |
| **pV** | 58.6 | N/A | 54.7 | N/A |
| **pVI** | 60.3 | 6.2 | 62.1 | 9.25 |
| **pVIII** | 78.6 | 1.99 | 79.8 | 1.13 |
| **pIX** | 55.6 | 1.81 | 77.2 | 7.83 |

**Supplementary Table 4:** Amino acid sequences of hypervariable regions (HVRs) of the HAdV-C5, -D26, -F40 and -F41.

| | HAdV-C5 | HAdV-D26 | HAdV-F410 | HAdV-F41 |
|---|---|---|---|---|
| **HVR1** | DEAATALEI NLEEEDDD NEDEVDEQ AEQQKTHV F | ETKEKQGTTGGVQ QEKDVTKTF | TNQNKTNSFGQ | KDNNKIKVRGQ |
| **HVR2** | VEGQTPK | TDETAENGKKDI | LDSNNRDV | TDTTNQPI |
| **HVR3** | YETEINH | QENEAF | NINPMQN | NSEVGAAQK |
| **HVR4** | GILVKQQN GKLESQ | AKFKPVNEGEQPK DLD | AKLVKNDDNQT TTTN | ASLITNGTDQTL TSD |
| **HVR5** | STTEATAGN GDNLTPK | DVPGGSPPAGGSGE EYKAD | TTATETANFSPK | ALPSTPNEPK |
| **HVR6** | TIKEGNSRE L | GTSDNSSEIN | DVNGTSAELL | DVAQGTISSADL |
| **HVR7** | GGVINTEEL EKVKPKTG QENGQWEK DATEFSDKN EIRVGNNF | BGTGTNSTYQGVKI TNGNDGAEESEWE KDDAISRQNQICKG NVY | NGQGUSNSYQG VKTDNGTNWSQ NNTDVSSNNEISI GNVF | GGSAATDTYSGI KANGQTWTAD DNYADRGAEIES GNIF |

**Supplementary Table 5:** Possible interactions formed between HAdV-F41 pV and hexon chains.

| | H2.1 | H3.2 | H3.3 | H4.1 | H4.2 |
|---|---|---|---|---|---|
| Gln170 | H-bond with carbonyl oxygen of Asn332 | H-bond with Ser61 | --- | --- | --- |
| Pro171 | --- | --- | --- | --- | --- |
| Thr172 | H-bond with either Asn707* or carbonyl oxygen of Ala920* | --- | --- | --- | --- |
| Met173 | --- | --- | --- | --- | --- |
| Gln174 | H-bond with carbonyl oxygen of Gly921 | --- | H-bond with His872 | --- | --- |
| Leu175 | --- | --- | --- | --- | --- |
| Met176 | --- | --- | --- | --- | --- |
| Val177 | --- | --- | --- | --- | --- |
| Pro178 | --- | --- | --- | --- | --- |
| Lys179 | --- | --- | Electrostatic interaction with Asn869 | --- | --- |
| Arg180 | Electrostatic interaction with Asn869* or Asp699* | --- | --- | --- | Electrostatic interaction with Ser642* |
| Gln181 | H-bond with carbonyl oxygen Tyr867 | --- | --- | --- | --- |
| Lys182 | --- | --- | --- | --- | Electrostatic interaction with Asn869 |
| Leu183 | --- | --- | --- | --- | --- |
| Glu184 | --- | --- | --- | --- | H-bond with Asn869 |
| Glu185 | --- | --- | H-bond with Asn644 | --- | --- |
| Val186 | --- | --- | --- | --- | --- |
| Leu187 | --- | --- | --- | --- | --- |
| Glu188 | --- | --- | Electrostatic interaction with Arg643. H-bond with Ser919 | --- | --- |
| Asn189 | --- | --- | --- | --- | --- |
| Met190 | --- | Van-der-Waals interaction formed with hydrophobic pocket formed by residues of H3.2 and H3.3. | Van-der-Waals interaction formed with hydrophobic pocket formed by residues of H3.2 and H3.3. Backbone interactions suggest inter-molecular β-sheet. | --- | --- |
| Lys191 | --- | --- | Backbone interactions suggest inter-molecular β-sheet. | --- | --- |
| Val192 | --- | --- | Backbone interactions suggest inter-molecular β-sheet. | --- | --- |
| Asp193 | --- | --- | Backbone interactions suggest inter-molecular β-sheet. | H-bond with Ser61 | --- |

**Supplementary Table 6:** Data collection and model building and refinement statistics.

| | HAdVF-41 pH 7.4 (1st dataset / 2nd dataset) | HAdVF-41 pH 4.0 | HAdVF-41 PB (tilted) | HAdVF-41 PB (untilted) |
|---|---|---|---|---|
|  | EMD-YYYYY / PDB XXXX | EMD-YYYYY | EMD-YYYYY / PDB XXXX | ----- |
| Data Collection and Processing | | | | |
| Magnification | 130,000 | 130,000 | 130,000 | 130,000 |
| Voltage (keV) | 300 | 300 | 300 | 3000 |
| Electron exposure ($e^-$/Å$^2$) | 40 | 40 | 40 | 40 |
| Detectore dose ($e^-$/Å$^2$/s) | 8.8 / 5.6 | 9.4 | 7.77 | 7.39 |
| Dose per frame ($e^-$/Å$^2$) | 1.03 / 1.08 | 2.4 | 0.93 | 1.11 |
| Defocus range (µm) | -0.8 to -2.5 | -0.7 to -2.7 | -0.5 to -3.0 | -1.0 to -2.5 |
| Pixel size (Å) | 1.041 | 1.041 | 1.041 | 1.041 |
| Stage tilt | --- | --- | 30° | --- |
| Initial particle number (total) | 7151 / 18,516 (25,667) | 6443 | 198,491 | 232,000 |
| Final particle number (total) | 4150 / 15,322 (19,472) | 4905 | 107,110 | 216,910 |
| Average map resolution (Å) | 3.77 | 4.96 | 3.73 | 3.26 |
| FSC threshold | 0.143 | 0.143 | 0.143 | 0.143 |
| Refinement and Model building | | | | |
| Initial Model used | 5TX1 | HAdVF-41 | HAdVF-41 (Chain M) | ---- |
| Model resolution (Å) | 4.0 | ----- | 3.85 | ---- |
| FSC threshold | 0.143 | ---- | 0.143 | ---- |
| Map sharpening range (Å$^2$) | -14.69 to 308.53 | ---- | 4.76 to 224.29 | ---- |
| Number of sub-maps for sharpening | 891 | ---- | 230 | ---- |
| Model composition | | | | |
| Non-hydrogen atoms | 94964 | --- | 16665 | --- |
| Protein residues | 11973 | --- | 2080 | --- |
| Number of chains | 36 | --- | 5 | --- |
| R. m. s. deviations | | | | |
| Bond lengths (Å) | 0.017 | --- | 0.006 | --- |
| Bond angles (°) | 2.436 | --- | 0.759 | --- |
| Validation | | | | |
| MolProbility score | 3.99 | --- | 2.65 | --- |
| Clashscore | 97.13 | --- | 20.74 | --- |
| Poor rotamers [%] | 8.53 | --- | 0.00 | --- |
| Map-to-model CC (masked) | 0.79 | --- | 0.85 | --- |
| Map Resolution (Model-vs-map) | 4.0 | --- | 4.0 | --- |
| FSC threshold | 0.5 | --- | 0.5 | --- |
| Ramachandran plot | | | | |
| Favoured (%) | 71.67 | --- | 70.69 | --- |
| Allowed (%) | 20.13 | --- | 28.57 | --- |
| Outliers (%) | 8.2 | --- | 0.74 | --- |

**Supplementary Table 7:** Data processing statistics of the localised asymmetric reconstruction of the HAdV-F41 penton base and HVR2-containing loop.

| | HAdVF-41 penton base | HAdVF-41 HVR2-containing loop |
|---|---|---|
| | EMD-YYYYY | EMD-YYYYY |
| Sub-particle identification, extraction and filtering | | |
| Copy number | 60 | 60 |
| Symmetry | I | I |
| Initial sub-particles number | 1,168,320 | 1,168,320 |
| Final sub-particles number | 412,038 | 467,190 |
| Sub-structure reconstruction | | |
| Symmetry imposed | C1 | C1 |
| FSC threshold | 0.143 | 0.143 |
| Average resolution | 2.99 | 3.35 |

*Supplementary Methods: Structural bioinformatics workflow to determine the identity of an unknown protein chain*

Well-resolved continuous electron density was found for a 24 residue protein chain at the interior of the HAdV-F41 capsid in between the three non-peripentonal hexons (Fig. 4A,B). The local resolution at this position was ~2.8 Å as estimated by ResMap (Supplementary Fig. 1). At this resolution, a direct sequence determination from the structure was not possible. We reasoned that the electron density might still contain sufficient information to reveal the identity of the unknown chain if combined with other information. To this end, a structural bioinformatics workflow was conceived (Supplementary Figure 7).

Manual model building in Coot(*59*) allowed for the placement and initial real-space refinement of a 24mer poly-alanine trace into the electron density. Inspection of the placed model revealed density corresponding to large side chains at several positions. This was used as an initial constraint to narrow the search space consisting of the 32 unique proteins of the HAdV-F41 proteome. Amongst the positions with density for larger side chains, we selected positions 4 and 21 in the chain for the initial search constraint. These positions carried less electron density than observed for any Trp residue in the remaining HAdV-F41 structure, but had electron density consistent with any other large side chain. Thus, a sequence search pattern was devised by fixing these positions to allow only such large side chains. During building of the HAdV-F41 model, Asp and Glu residues that were neither spatially constrained nor formed at least one interaction with neighbouring residues were not observed to have electron density covering their side chain carboxy group. Since the side chains at positions 4 and 21 in the chain are not in a chemical surrounding that would allow for such interactions, Asp and Glu were excluded from the search. Positions 4 and 21 were thus allowed to be either of the residues (R,K,M,N,Q,F,Y). With this constraint the sequence of all HAdV-F41 proteins were scanned(*67*), allowing for either N-to-C ("forward") or C-to-N ("reverse") direction of the peptide. This search resulted in 942 candidate sequences (Supplementary Data 1).

Next, we scored the candidate sequences according to how well they fit the experimental electron density. To that end, the following steps were performed for candidate sequences of both "forward" and "reverse" direction: (1) the poly-alanine trace was mutated to each identified sequence in Coot(*59*) followed by (2) real-space refinement of the model against the experimental density in Phenix(*58*). To ensure proper behaviour, real-space refinement was performed in the presence of the protein chains immediately surrounding the peptide model. Finally, (3) per-chain map-to-model correlation coefficients ($CC_{map\text{-}to\text{-}model}$) were calculated in Phenix as a means to evaluate their fit to the electron density. All 942 chains were ordered according to their $CC_{map\text{-}to\text{-}model}$. When plotting the values separately for each direction of the polypeptide chain, the "forward" direction had consistently higher $CC_{map\text{-}to\text{-}model}$ values than the "reverse" direction (Supplementary Figure 7). Inspection of several fitted models of the "reverse" direction also made it clear that this direction is stereochemically incompatible with the electron density. Thus, we proceeded only with the candidate sequences from the "forward" list. Visual inspection of several fitted "forward" models in the electron density map made clear that the higher scoring sequences had a better fit to the map. Below a $CC_{map\text{-}to\text{-}model}$ value of 0.76 models started to display several major inconsistencies with the experimental electron density. Thus, all sequences of the "forward" direction with a $CC_{map\text{-}to\text{-}model}$ $\geq 0.76$ were taken forward (Supplementary Data 1).

Next, we incorporated information from the proteomics analysis of the purified HAdV-F41 virions. The proteomic analysis was performed using a filter assisted sample preparation (FASP)(*31*), which allows a more complete recovery of samples compared to gel extractions followed by in-gel digestions, and it notably identified all known capsid proteins along with several other proteins of the HAdV-F41 proteome. Excluding candidates not present in this proteome lead to the exclusion of a small number of candidates, e.g. some "early" adenovirus proteins (Supplementary Data 1).

The remaining list of 27 sequences was manually curated to remove twelve sequences belonging to the fibre proteins, and built parts of the hexon, penton base and protein IIIa - i.e. sequences that are clearly established to reside in different tertiary structure in another location in the virion. After the steps described above, a final list of 15 candidates remained (Supplementary Methods Table 1, Supplementary Data 1). Of these 15 candidates, seven were disqualified after thorough visual inspection of model fit to map since the model had a small side chain (Gly or Ala) with no possibilities for chemical modification at a position in the map where there was clear presence of additional side chain electron density (seq351, seq374, seq521, seq544, seq564, seq729, seq763). Of the two remaining E2A DBP candidate sequences, known to be a subunit of the adenoviral polymerase complex, one (seq541) was excluded based on being part of a β-sheet and a Zn-binding motif in the crystal structure of HAdV-C5 (PDB: 1ANV)(*68*). The other E2A DBP sequence (seq520) as well another three sequences (seq81, seq352, seq348) were excluded since they all had proline-containing motifs which could not be modelled into the experimental electron density without violating geometrical restraints (Supplementary Methods Table 1).

After this, three candidates remained. Of these three, two belonged to protein IIIa (seq234 and seq235) and overlapped in eight positions in their respective sequences. These sequences both reside in the C-terminal ~1/3 of pIIIa, which had not been built in the HAdV-F41 model. These pIIIa sequences were excluded based on the following reasoning: Firstly, it must be noted that there was sufficient unbuilt sequence of pIIIa that it could theoretically stretch across from where its N-terminal 2/3 are located, near the five-fold symmetry axis of the penton base, to the unknown chain.

However, we noticed that the published structure of HAdV-D26(*21*) had clear electron density and a built model for virtually all of pIIIa, including the sequences corresponding to our candidate sequences. In HAdV-D26, these sequences are present in a small, tightly folded α-helical domain with high sequence identity to the HAdV-F41 pIIIa (Supplementary Table 2, Supplementary Figure 7), crucially including a 80% sequence identity for the sequence that both candidate sequences span. We used SWISS-MODEL(*56*) to construct a homology model for HAdV-F41 pIIIa, including the part not built in our structure (Supplementary Figure 7). The algorithm identified HAdV-D26 pIIIa as the closest hit and built a homology model based on this protein. Crucially, the per-residue local quality estimate of the homology model is equally high in the part of pIIIa which was built in the HAdV-F41 structure (and which we know to be similar to the homology model) and in the domain containing the candidate sequences. The lowest per-residue model quality, on the other hand, was ound in a linker region connecting the part that we could build in the HAdV-F41 structure and the part containing the pIIIa candidate peptide (Supplementary Figure 7). Based on the above considerations, we considered it highly likely that the pIIIa candidate sequences are present in a domain of similar fold to the domain built in HAdV-D26 pIIIa, and therefore not to be the unknown chain. The only remaining candidate sequence belongs to the core-capsid bridging protein V, which has not yet been identified in any published adenovirus structure. Thus, by sequential exclusion of candidates, we arrived at pV as the most likely candidate for the unknown chain.

Supplementary Methods Table 1: List of final candidates.

| Model | Sequence | $CC_{m2m}$ | ProteinID | Comments |
|---|---|---|---|---|
| seq348 | VLENMKVDPSVEPEVKVRPIKEIG | 0.7765 | V | Arg18 pushed out N-terminus of neighbouring hexon. After manual correction and re-evaluation the CCm2m was 0.7623. All three proline residus are located outside of the density and cannot be corrected without violating geometrical restraints. |
| seq541 | SVVQIKNDDARCCAEDVSCGNNMF | 0.7726 | E2A DBP | Known location in β-sheet and Zn-binding motif of HAdV-C5 DBP (PDB: 1ANV). |
| seq374 | SGIKNFGSSIKSFGNKAWNSNTGQ | 0.7708 | VI | Two Glycine in position where there is additional density |
| seq729 | LPQQERAGAEENNEGERQSAQEPL | 0.7702 | E2B DNA pol | Two Glycine in position where there is additional density |
| seq763 | VGHNINGFDEIVLAAQVINNRSDV | 0.7685 | E2B DNA pol | Two Glycine in position where there is additional density |
| seq520 | VIPRDLTPPEEEENNQSGSSKAVT | 0.7677 | E2A DBP | Double proline motif is located outside of density and cannot be corrected without violating geometrical restraints. |
| seq521 | EEENNQSGSSKAVTMLITNPQVDP | 0.7673 | E2A DBP | Glycine in position where there is additional density |
| seq346 | QPTMQLMVPKRQKLEEVLENMKVD | 0.7673 | V | Glutamic acid in a position with large side chain density however highly coordinated, suggesting stabilisation of carboxy-group. |
| seq544 | SCGNNMFSSKSCGLFFSEGLKAQI | 0.7651 | E2A DBP | Glycine in position where there is additional density. |
| seq235 | TARNMEPSLYSSNRPFINRLMDYL | 0.7649 | IIIa | Homology to pIIIa in HAdV-D26 predicts different location |
| seq352 | QTDQPAAVTTREIGLQTDPRYESV | 0.7648 | V | Two proline residues are located outside of density and cannot be corrected without violating geometrical restraints. |
| seq351 | AVAMAEAMETQTDQPAAVTTREIG | 0.7644 | V | Alanine in position where there is additional density. |
| seq564 | ANLFSPRVPPKKQANGTCEPNPRL | 0.7638 | 100k | Glycine in position where there is additional density. |
| seq234 | LYLMREGMTPSSALDMTARNMEPS | 0.7638 | IIIa | Homology to pIIIa in HAdV-D26 predicts different location |
| seq81 | SMMNEVTPLLREDGSCSSLNYQLQ | 0.7612 | IVa2 | Threonine and proline residue located outside of density and cannot be corrected without violating geometrical restraints. |

**Supplementary Materials not included in this manuscript file**

**Supplementary Data 1 (spreadsheet):**

List of potential unknown chain candidates. Each individual spreadsheet tab represents a filtering step.

**Supplementary Movie 1:**

Structure of HAdV-F41 at pH=7.4, shown as a grey surface. The asymmetric unit (ASU), used for model building, is highlighted in red. Final state shows the built model of the HAdV-F41 ASU in cartoon representation in the volume of the ASU, shown as a semi-transparent grey surface.

**Supplementary Movie 2:**

The structure of the HAdV-F41 asymmetric unit shown in cartoon representation and surface representation. The color scheme is identical to the scheme in Fig. 1.

**Supplementary Movie 3:**

Assembly-induced conformational changes of the HAdV-F41 penton base (PB). The morph movie shows the changes of the PB between the free and virus bound state. The PB is shown in cartoon representation with individual chain of the homopentamer colored differently.

**Supplementary Movie 4:**

The DNA-binding protein V binds to the HAdV-F41 at a conserved site. The pV density is shown as a maroon surface. The final placed model is shown in stick representation placed in a grey semi-transparent surface.
